# Engineering human protein switches for functional control of CARs and transcription factors via oral drug administration

**DOI:** 10.1101/2025.10.17.683068

**Authors:** Elise Sylvander, Álvaro Muñoz-López, Giulia D’Accardio, Dominik Emminger, Konstantina Mouratidis, Fabian Engert, Hayeon Baik, Sanjana Balaji, Theresa Michls, Michelle C. Buri, Daniel Maresch, Anna Urbanetz, Markus Schäfer, Joerg Mittelstaet, Johannes Zuber, Antonio Rosato, Boris Engels, Charlotte U. Zajc, Michael W. Traxlmayr, Manfred Lehner

## Abstract

While CAR T cells have revolutionized the treatment of certain hematologic malignancies, they can cause severe toxicities, which are expected to be exacerbated with next-generation CAR Ts engineered for improved proliferation, persistence, and efficacy. Therefore, regulatory systems are urgently needed to be able to control these living drugs directly in patients. Here, we engineered a molecular switch, in which the interaction of two human proteins is efficiently induced with the orally available and non-toxic drug A1120. We demonstrate the versatility of this switch by regulating CAR signaling and transcriptional activity in human T cells *in vitro* and *in vivo*. Both systems were tightly controlled in the absence of the drug but strongly activated upon administration of the small molecule. Since this switch enables the regulation of diverse systems including CARs and transcription factors, we anticipate that it represents an important step towards next-generation cellular therapies with improved safety and efficacy.

## Introduction

CAR T cells have revolutionized the treatment of diverse hematologic malignancies including relapsed/refractory B cell leukemias and lymphomas, as well as multiple myeloma^1^. However, in a large fraction of patients, the high potency of CAR T cells also translates to severe toxicities, including cytokine release syndrome (CRS), immune effector cell-associated neurotoxicity syndrome (ICANS) and on-target/off-tumor toxicity, the latter being particularly problematic with solid tumors^2^. While CRS can be clinically managed with corticosteroids or tocilizumab, patients experiencing ICANS do not always respond and on-target/off-tumor toxicities represent an even greater problem, which – depending on the antigen – has even resulted in fatalities^3^. Importantly, toxicities are expected to be even more frequent and severe with next-generation CAR T cells engineered for improved potency, proliferation and persistence^4–10^. Unfortunately, discontinuing treatment is not an available option for CAR T cells, since these living drugs proliferate after administration to the patient and can persist for more than a decade^11^.

Thus, to address this major limitation of CAR T cells, regulatable systems are urgently needed. One option is the co-expression of suicide switches, which enable drug-induced elimination of the CAR T cells. However, while this concept has been shown to rapidly resolve toxicities^12^, it leads to irreversible ablation of the expensive and potentially life-saving therapy. Another concept is the use of drug-regulated switches, in which protein dimerization is induced upon drug administration (also known as chemically induced dimerization, CID). The key advantages of such switches are their reversibility, i.e. the ability to transiently deactivate a given signaling component, as well as their versatility and modularity, enabling the regulation of diverse cellular functions^9,13–17^. Although much progress has been made in the development of drug-responsive switches, the clinical translation of these systems is limited due to the use of non-human protein components and/or side effects of the small molecule drugs^9,13–15,18–25^.

For example, we previously developed a switch system based on human retinol binding protein 4 (hRBP4), which undergoes a conformational change upon loading with the small molecule drug A1120^13,26,27^. This drug-induced conformational change can be specifically recognized by an engineered binder, thus resulting in A1120-dependent protein dimerization (Fig. 1A). The major advantages of this switch system include the use of an orally available and non-toxic drug^26,28,29^, the human origin of hRBP4, as well as the flexibility of the platform, which enabled us to use two different scaffolds for binder engineering: reduced-charge Sso7d (rcSso7d)^30^ and the 10^th^ type III domain of human fibronectin (Fn3)^31^. However, in its prototype version, this switch system also came with several disadvantages: (i) due to the presence of three disulfide bonds in hRBP4, this protein is expected to be non-functional in the reducing environment of the cytoplasm, thus limiting its applicability to the extracellular space; (ii) A1120 binds to endogenous hRBP4 present in human plasma with identical affinity; (iii) both engineered binders suffer from certain limitations: RS3 is derived from the non-human scaffold rcSso7d, thus raising immunogenicity concerns, whereas RF2 derived from the human Fn3 domain only showed limited switching behavior (only 16-fold increase in affinity in the presence vs. absence of A1120) and is relatively unstable with a *T*_m_ of only 48 °C^13^.

**Figure 1.**
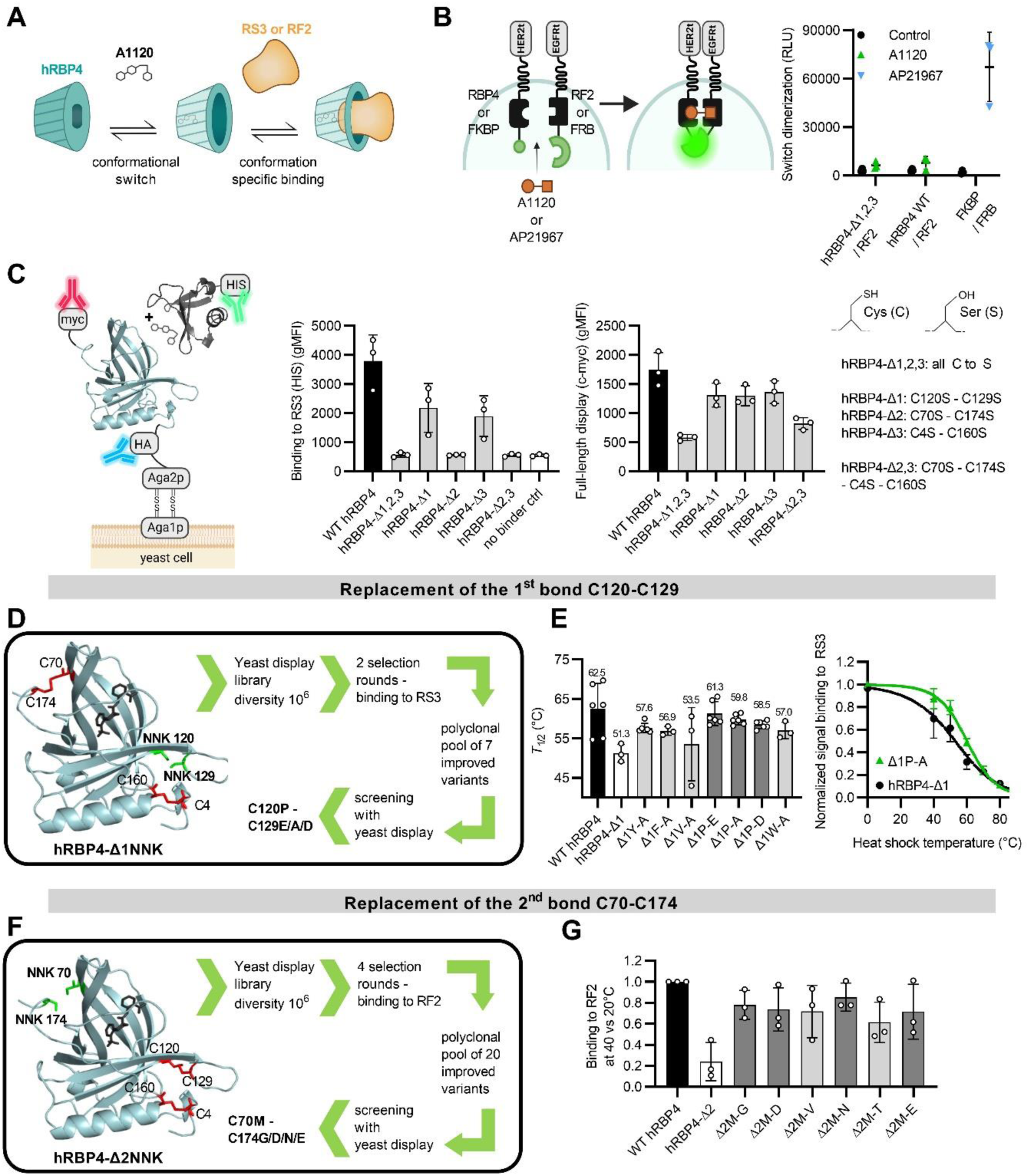
Testing of intracellular functionality of WT-hRBP4-based switches and engineering of the first two disulfide bonds. **A)** Principle of the hRBP4-based switch. The binders RS3 and RF2 specifically recognize the A1120-loaded conformation of the lipocalin hRBP4, human retinol binding protein 4; RS3 is derived from the Sso7d protein of *Sulfolobus solfataricus* and RF2 from human fibronectin type III (FN3) domain. **B)** Analysis of A1120-induced dimerization of cytoplasmic RBP4 variants and RF2 using the NanoBiT® system. A1120 (5 µM) or AP21967 (0.5 µM) was added as indicated for dimerization of hRBP4 / RF2 or FKBP / FRB, respectively. hRBP4-Δ1,2,3 has all 6 Cys residues mutated to Ser. RLU: relative light units. **C)** Representation of hRBP4 displayed on yeast binding to soluble RS3 (left), as well as RS3 binding in the presence of 5 µM A1120 (middle) and expression (right) of hRBP4 mutants with deleted disulfide bonds. gMFI: geometric mean fluorescence intensity (mean ± SD, n=3). **D)** Schematic overview of the engineering process for Δ1 C120 - C129. **(E)** *T*_1/2_ values of hRBP4-Δ1NNK mutants. Binding to RS3 was used to monitor hRBP4 unfolding in heat exposed yeast cells (in the presence of 300 nM RS3 and 5 µM A1120). The average *T*_1/2_ for each mutant is indicated on the left (mean ± SD, n=3-6), and representative denaturation curves are shown on the right. **F)** Schematic overview of the engineering process for Δ2 C70 - C174. **(G)** Binding of the top 7 hRBP4-Δ2NNK mutants to RF2 measured at 40 °C versus 20 °C in the presence of 5 µM A1120. This ratio was normalized to the one of the WT hRBP4 (mean ± SD, n=3).

To address these limitations, we have now optimized both protein components through extensive protein engineering. Briefly, we generated disulfide-free hRBP4 versions which enabled intracellular functionality, we increased their affinity to A1120 and optimized the human-derived binder RF2 by stability engineering and by strongly improving its A1120-dependent switching behavior. Furthermore, we used the optimized switch system to regulate both CARs and transcription factors *in vitro* and *in vivo*, demonstrating its versatility and efficiency in controlling cell function.

## Results

### Switches based on wild type hRBP4 are dysfunctional in the cytoplasm of T cells

We hypothesized that the reducing intracellular environment would impair formation of the three disulfide bonds in wild type (WT)-hRBP4, thus leading to dysfunctional switches. To test this, we assessed switches based on either WT-hRBP4 or a variant thereof lacking all three disulfide bonds (hRBP4-Δ1,2,3; all Cys mutated to Ser) in the cytoplasm of Jurkat T cells. In contrast to the benchmark switch (FKBP/FRB), neither WT-hRBP4 nor hRBP4-Δ1,2,3-based switches were functional (Fig. 1B). Furthermore, when displayed extracellularly on yeast, WT-hRBP4 showed efficient expression and switch function, which was lost upon gradual deletion of individual disulfide bonds (Fig. 1C). Overall, these data demonstrate the dependency of hRBP4 on its disulfide bonds, leading to loss of function in the reducing intracellular environment.

### Engineering disulfide-free hRBP4 (dfRBP4) variants to enable intracellular function

To eliminate endogenous disulfide bonds in WT-hRBP4, we screened for optimal amino acid replacements at those positions by using yeast surface display technology. The first disulfide bond (C120-C129) was deleted by randomly mutating these positions with NNK codons and screening the resulting library (hRBP4-Δ1NNK) for maintained binding to RS3 in the presence of the drug A1120 (Fig. 1D). This yielded several amino acid combinations (Extended Data Fig. 1A) that enable efficient switch function and proper expression (Extended Data Fig. 1B and 1C). Of note, the three top candidates Δ1P-E, Δ1P-A and Δ1P-D are considerably more stable than the variant hRBP4-Δ1 (containing Ser mutations) and almost reach the stability of WT-hRBP4 despite the loss of one disulfide bond (Fig. 1E).

The second disulfide bond (C70-C174) was replaced with a similar approach. However, since RS3 interacts with C70 (Extended Data Fig. 1D) and therefore binding is lost upon mutation of this disulfide bond, we instead used binder RF2. Again, multiple amino acid combinations were found to be enriched (Extended Data Fig. 1E) and the top candidates (Δ2M-G, Δ2M-D, Δ2M-N and Δ2M-E) showed strong A1120-dependent binding to RF2, high expression levels (Extended Data Fig. 1F and 1G) and improved thermostability compared to the Ser-containing variant hRBP4-Δ2 (Fig. 1G).

Finally, to obtain entirely disulfide-free hRBP4 (dfRBP4) variants, we generated a library comprising the top-performing amino acid combinations at the first two disulfide bonds (P-E, P-A and P-D for C120-C129; M-G, M-D, M-N and M-E for C70-C174), as well as fully randomized positions of the last disulfide bond (C4-C160). Screening for A1120-dependent RF2 binding led to a mediocre dfRBP4 variant (V29) with limited binding to RF2 and poor expression level (Extended Data Fig. 2A, 2B and 2C). However, both parameters could be improved by introducing additional diversity via error-prone PCR (epPCR), yielding variants V1 to V28. To further enhance expression levels and protein folding, we added a final round of epPCR and screened for efficient full-length protein expression (variants V30 to V39). Systematic comparison of all variants (Extended Data Fig. 2B, 2C, 2D and 2E) revealed a set of 14 top candidates, which were subsequently analyzed in detail with respect to RF2 binding (Extended Data Fig. 3A), expression on yeast (Extended Data Fig. 3B), affinity to RF2 in the presence of A1120 (Extended Data Fig. 3C), aggregation of solubly expressed proteins (Extended Data Fig. 3D) and switch function in the cytoplasm of Jurkat T cells (Extended Data Fig. 3E). One notable finding was the superiority of the variants V30 to V39, in particular with respect to protein aggregation (Extended Data Fig. 3D), demonstrating the importance of the additional final engineering round that yielded these variants. Based on all these data, we chose the dfRBP4 variants V30, V33 and V35 for further experiments.

### Detailed biochemical analysis of optimized dfRBP4 variants

Despite the elimination of all three disulfide bonds, the dfRBP4 variants V30, V33 and V35 are highly stable with *T*_m_ values above 60 °C and efficiently expressed in the cytoplasm of primary human T cells (Fig. 2F and 2G). Moreover, they show improved affinity and binding signal to RF2 compared with WT-hRBP4 (Fig. 2B and Extended Data Fig. 3A) and similar extracellular expression levels (Extended Data Fig. 3B). Importantly, the engineering process also strongly decreased the *EC*_50_ of A1120 (13- to 112-fold, Fig. 2C). That is, when applied in patients, A1120 will preferentially bind to these engineered dfRBP4 variants compared to WT-hRBP4 circulating in the plasma.

**Figure 2.**
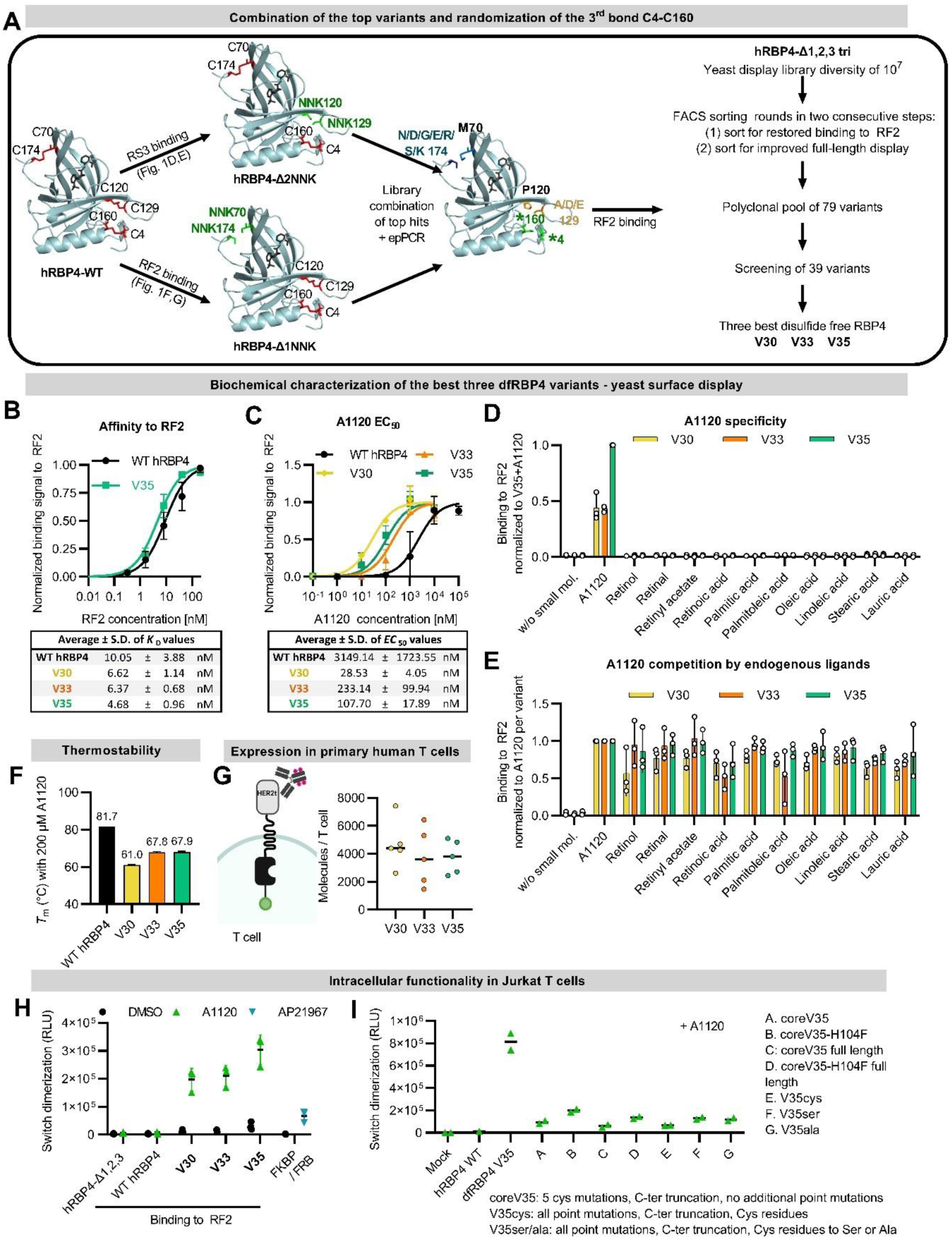
dfRBP4 engineering and characterization. **A)** Overview of the stabilization strategies. The library “hRBP4-Δ1,2,3 tri” included the mutations C120P and C70M, and was randomized at positions 129 (GMW), 174 (RRW), and at positions 4 and 160 to all amino acids except cysteine. Additional mutations were added by error prone PCR. The library was selected for binding to RF2 and later for improved full-length expression on yeast. **B)** *K*_D_ values were measured by titrating soluble RF2 on yeast displayed dfRBP4 variants in the presence of 5 µM A1120 (mean ± SD, n=3). The fitted curves for WT hRBP4 and dfRBP4 V35 are shown on top and all *K*_D_ values are indicated on the bottom. **C)** Binding of the yeast-displayed hRBP4 mutants to 200 nM soluble RF2 was measured in the presence of increasing concentrations of A1120 (mean ± SD, n=4). The fitted curves for all variants are shown on top and all *EC*_50_ values are indicated on the bottom. **D)** Binding to 100 nM soluble RF2 was assessed in the presence of 1 µM A1120 or 5 µM retinoid or 100 µM fatty acid or in the absence of any small molecule. Binding levels (gMFI) were normalized to binding of dfRBP4 V35 in the presence of 1 µM A1120 (mean ± SD, n=3). **E)** 100 nM soluble RF2 were incubated with dfRBP4 displayed on yeasts in the presence of 1 µM A1120 alone or in combination with 5 µM retinoids or 100 µM fatty acids as indicated. Loss of binding signal, indicating competition with A1120, was measured. Binding levels (gMFI) were normalized to the A1120 only condition (mean ± SD, n=3). **F)** DSC-determined *T*_m_ values of proteins loaded with 200 µM A1120 (mean ± SD, n=3). **G)** Expression of the constructs in primary human T cells. **H)** Analysis of A1120-induced dimerization of cytoplasmic RF2 and hRBP4 variants using the NanoBiT® assay. A1120 (5 µM) was added as indicated. Binding of FKBP to FRB in the presence of rapalog (AP21967, 0.5 µM) served as a benchmark (mean ± SD, n=3). RLU: relative light units. **I)** Analysis of the impact of individual mutations in dfRBP4 V35 on its intracellular functionality using the NanoBiT® assay. Dimerization of cytoplasmic RF2 and dfRBP4 V35 versions containing different mutations was measured in the presence of 5 µM A1120 (n=2).

Of note, only A1120, but none of the endogenous small molecules (various fatty acids and retinoids) known to bind to WT-hRBP4, was able to trigger these V30-, V33- or V35-based switches (Fig. 2D), demonstrating their pronounced specificity for A1120. Moreover, we investigated whether these endogenous compounds applied at physiological concentrations can compete with A1120 for dfRBP4 binding and thereby block switch function. Even though these endogenous small molecules were in 5-fold (retinoids) and 100-fold (fatty acids) excess compared with A1120, none of them efficiently competed with A1120, in particular in the V35-based switch system (Fig. 2E).

Since the primary goal of this study was the adaptation of hRBP4 for intracellular function, we now investigated whether the engineering of stable and disulfide-free RBP4 versions enabled potent functionality in the cytoplasm of human T cells. Remarkably, in contrast to WT-hRBP4 and the Ser-variant hRBP4-Δ1,2,3, efficient A1120-dependent switch function was obtained with V30, V33 and V35, even reaching higher levels of activation than the FKBP/FRB benchmark switch (Fig. 2H and Extended Data Fig. 3E), demonstrating the success of the engineering process.

Finally, we chose V35 as a representative candidate to analyze the impact of the respective mutations on its switch function (Fig. 2I). V35 contains 5 Cys substitutions, a C-terminal truncation of 7-amino acids, as well as three additional point mutations. We designed multiple variants containing different sets of mutations of V35: coreV35 (5 Cys substitutions and truncation), coreV35-H104F (includes H104F, since H104 is mutated in most enriched variants), full length versions of both, as well as V35 versions including all additional mutations, but in which the original Cys positions were either mutated back to Cys (V35 cys), Ser (V35ser) or Ala (V35ala). Of note, neither the Cys replacements alone (coreV35 and coreV35 full length) nor the additional mutations (V35cys) were sufficient for high functionality. Moreover, the optimal amino acid combinations that were enriched at the Cys positions clearly outperformed simple mutations to Ser or Ala (V35 vs. V35ser or V35ala) (Fig. 2I).

Together, these data clearly demonstrate that the complete set of V35 mutations is required for efficient switch function. Moreover, we demonstrated that switches based on these dfRBP4 variants are stable, well-expressed, show high specificity and improved sensitivity to A1120, as well as high functionality in the cytoplasm of T cells.

### Stability engineering of the second switch component RF2

After optimization of hRBP4, we turned our attention to the other protein component of the switch, i.e. the engineered binder RF2 based on the human Fn3 domain. We hypothesized that RF2 is suboptimal for CAR T cell applications due to its low thermostability with a *T*_m_ that is only slightly above body temperature (48 °C^13^).

Therefore, we sought to stabilize RF2 by screening a randomly mutated RF2 library for maintained binding to a conformationally specific ligand (V35 loaded with A1120) after heat incubation, i.e. for resistance to thermal denaturation. A1120-loaded V35 was chosen as the detection reagent for correctly folded RF2, since it yielded the highest binding signal among all dfRBP4 variants in a preliminary experiment (Fig. 3A). After multiple rounds of selection (Fig. 3B), we obtained seven RF2 mutants, all of which showed strongly increased thermal stability on yeast (Fig. 3C, Extended Data Fig. 4A and 4B). This improved thermostability was subsequently confirmed in a soluble expression format for the top five candidates (*T*_m_ of all variants increased by >10 °C; Fig. 3D), showing a strong correlation with the *T*_1/2_ determined in the yeast display format (Extended Data Fig. 4C). These soluble proteins were further analyzed by SEC-HPLC, revealing largely monomeric proteins resistant to aggregation (Fig. 3E). RF2 shows delayed elution from the SEC column when compared with the Fn3-WT domain, indicating some weak interactions with the column matrix, which is frequently observed with engineered proteins^30,32–34^. Of note, this elevated retention on the column was largely reverted in the stabilized variant RF2s11, presumably due to reduced hydrophobicity on its surface (Fig. 3E, Extended Data Fig. 4A).

**Figure 3.**
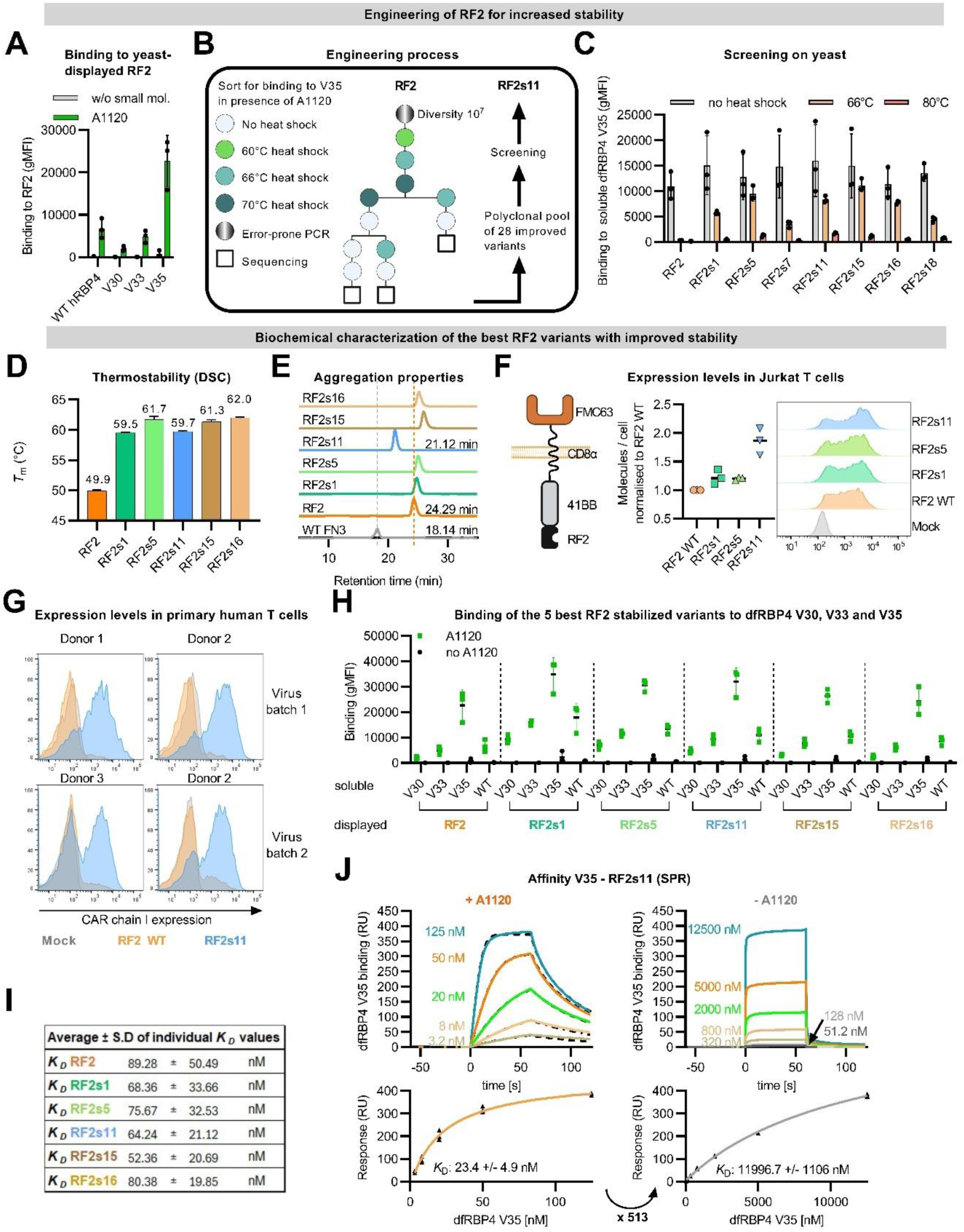
Engineering of RF2 for increased stability. **(A)** Binding of 200 nM soluble disulfide-free hRBP4 variants to yeast displayed RF2, assessed in the presence or absence of 5 µM A1120 (mean ± SD, n=3). **(B)** Schematic overview of the engineering process for RF2 stabilization. **(C)** Binding of yeast-displayed RF2 and derived stabilized mutants to 20 nM soluble dfRBP4 V35 after a 66 °C or 80 °C heat shock, or without heat shock. Binding was assessed in the presence of 5 µM A1120 (mean ± SD, n=3). **(D)** DSC-determined *T*_m_ values of RF2 and derived stabilized mutants (mean ± SD, n=3). **(E)** Size exclusion chromatography profile of RF2 variants. One representative measurement of three independent experiments is shown. The WT fibronectin domain (WT FN3), from which RF2 is derived, was included as a control. **F)** Expression of RF2 (stabilized/non-stabilized) in Jurkat cells. Jurkat cells were electroporated with mRNA encoding the CAR constructs depicted on the left. The number of molecules per cell was quantified with flow cytometry 8 h after electroporation and normalized to the number of the CAR with the non-stabilized RF2 (mean ± SD, n=3). **G)** Expression of RF2 (stabilized/non-stabilized) in primary T cells. The same constructs as in (F) were stably expressed in primary human T cells via lentiviral transductions. Two different virus batches were used across 3 different T cell donors, and expression was assessed 13 days after transduction (n=4). **H)** Binding of soluble hRBP4 proteins (200 nM) to yeast-displayed RF2 variants, assessed in presence or absence of 5 µM A1120 (mean ± SD, n=3). **I)** *K*_D_ values measured by titrating soluble dfRBP4 V35 on yeast displayed RF2 variants in the presence of 5 µM A1120 (mean ± SD, n=3). **J)** Multi-cycle kinetic (MCK) surface plasmon resonance (SPR) experiments with dfRBP4 V35 titrated onto RF2s11 immobilized on a sensor chip. The left and right panels show binding of V35 to RF2s11 in the presence and absence of 5 µM A1120, respectively. Representative diagrams of three independent experiments are shown at the top. *K*_D_ values were calculated by steady-state analysis and are shown at the bottom. RU: response units.

Importantly, these improved surface properties of RF2s11, together with its increased stability, led to elevated expression levels in a CAR format on Jurkat T cells (Fig. 3F), which was even more pronounced in primary human T cells (Fig. 3G). Finally, we confirmed that switch function was maintained during stability engineering, as demonstrated by drug-dependent binding to all dfRBP4 mutants and similar affinities to V35 (Fig. 3H and 3I).

Based on all data described above, we chose V35 and RF2s11 as the most promising switch components and therefore analyzed this V35-RF2s11 switch in more detail by SPR (Fig. 3J). In the presence of the drug A1120, the proteins V35 and RF2s11 interact with each other with a high affinity (*K*_D_ of 23 nM), whereas this affinity is strongly reduced (*K*_D_ 11996 nM) in the absence of A1120. This corresponds to a 513-fold affinity increase upon addition of A1120 (Fig. 3J), which is a significant improvement compared with the original WT-hRBP4-RF2 switch that only shows a 16-fold affinity difference^13^.

Summing up, we successfully addressed all limitations of the original WT-hRBP4-RF2 switch: (i) we engineered dfRBP4 variants to enable intracellular functionality in the cytoplasm of T cells, (ii) we improved the affinity to A1120, resulting in preferential A1120 binding to the switch as compared with endogenous WT-hRBP4, (iii) we stabilized the human binder RF2 and thereby elevated its expression in a CAR format in human T cells and (iv) we strongly improved the A1120-dependency of the switch (affinity of the two protein components in the presence vs. absence of A1120) from 16-fold to 513-fold.

### Drug-regulated CAR signaling using dfRBP4-based switches

Building upon this promising data, we integrated RF2s11 and either V35 or V33 into multiple switchable split CAR formats. A common feature of all these designs is the separation of the CD19-targeting scFv and the CD3ζ signaling domain onto two distinct polypeptide chains (chain I and II, respectively). However, the designs differ in their use of monomeric versus dimeric CAR chains, as well as the orientations of the switch components, i.e. which of them is used in which chain (Fig. 4A). To be able to measure CAR signaling with high sensitivity, we employed a previously validated Jurkat Nur77 reporter cell line^33^. This comprehensive experiment yielded several key insights: (i) most split CAR constructs showed efficient switch function, (ii) CAR signaling was strictly antigen-dependent, demonstrating absence of tonic signaling, (iii) V35 overall performs better than V33, (iv) dimeric chains generally produced stronger CAR signaling than their monomeric counterparts and (v) most dfRBP4-based split CARs outperformed the FKBP/FRB-based benchmark CAR^14^ (Fig. 4A).

**Figure 4.**
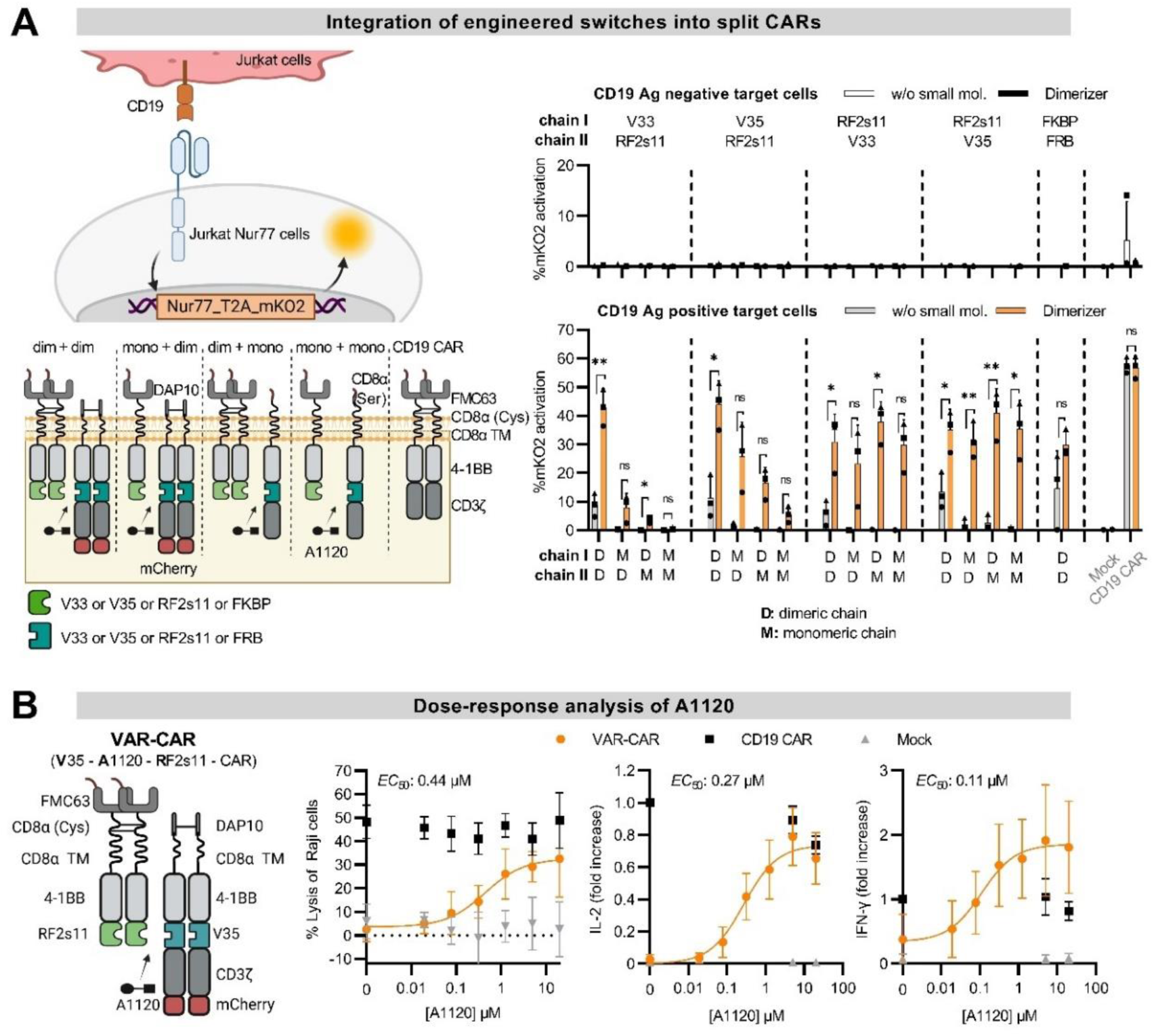
Drug-mediated regulation of split CAR function *in vitro*. **(A)** Integration of the switches into split CARs. A schematic of the Nur77 assay and the different split CAR architectures is shown on the left. Jurkat T cells, either electroporated or not with mRNA encoding the CD19 antigen, served as target cells and were co-cultured with Nur77 Jurkat reporter cells electroporated with split CAR constructs comprising the indicated on-switch. The appropriate dimerizer (5 µM A1120 for the RBP4 / RF2 and 0.5 µM rapalog for FKBP / FRB on-switch) was added as indicated. The order of the domains in the split CARs is indicated above the graph, and the dimerization status of each chain is shown below (D = dimeric, M = monomeric). mKO2 signal was measured by flow cytometry and Mock Nur77 Jurkat reporter cells (electroporated without mRNA) served as control (mean ± SD, n=3). **(B)** Dose-response analysis of A1120. Primary human T cells were transduced with either a standard CD19 CAR or the split CAR depicted on the left, and co-cultured with Raji target cells in the presence of increasing concentrations of A1120 (E:T of 2:1, 24 h). Lysis of the Raji cells (left, n=5, 2 different T cell donors) was normalized to lysis by Mock T cells without A1120. Secretion of IL-2 (middle) and IFN-γ (right) was normalized to that of the CD19 CAR without A1120 (n = 6, 3 different T cell donors).

To confirm these observations in a more physiological system, we tested all V35-based split CARs in primary human T cells and observed similar effects when measuring IFN-γ and IL-2 secretion, as well as cytotoxicity against two different target cell lines (Extended Data Fig. 5A).

Based on this comprehensive comparison of different split CAR architectures, we chose a final split CAR format with a dimeric RF2s11-containing chain I and dimeric V35-based chain II, which we term **V**35-**A**1120-**R**F2s11-CAR (VAR-CAR, Fig. 4B). Co-culture assays with primary human VAR-CAR T cells and Raji target cells yielded EC_50_ values in the sub-µM range, demonstrating the high sensitivity of the VAR-CAR (Fig. 4B). Furthermore, kinetics experiments demonstrated that the VAR-CAR is rapidly turned on and off upon A1120 addition and removal, respectively (Extended Data Fig. 5B).

### Control of VAR-CAR T cell function *in vivo* via oral drug administration

Next, we tested whether the anti-tumor activity of VAR-CAR T cells can be regulated *in vivo* by administering A1120 via the food in a Raji lymphoma mouse model (Fig. 5A). Of note, delivery of A1120 to the mice via the chow did not cause any toxicities (Extended Data Fig. 6A) and resulted in a median plasma concentration of 10.3 µM (Fig. 5C). In line with our *in vitro* data, showing full activation of VAR-CAR T cells in this concentration range (Fig. 4B), VAR-CAR T cells were highly active *in vivo*. Although slightly less effective than a conventional direct CD19-CAR, the split CAR design in the presence of A1120 enabled potent tumor control (Fig. 5B). In contrast, feeding mice with the same chow not containing A1120 completely abrogated anti-tumor activity of VAR-CAR T cells, as evidenced by identical tumor outgrowth in mice treated with VAR-CAR T cells and Mock T cell controls. These data underscore the tight control of the VAR-CAR activity *in vivo* in the absence of drug-induced VAR-CAR assembly.

**Figure 5.**
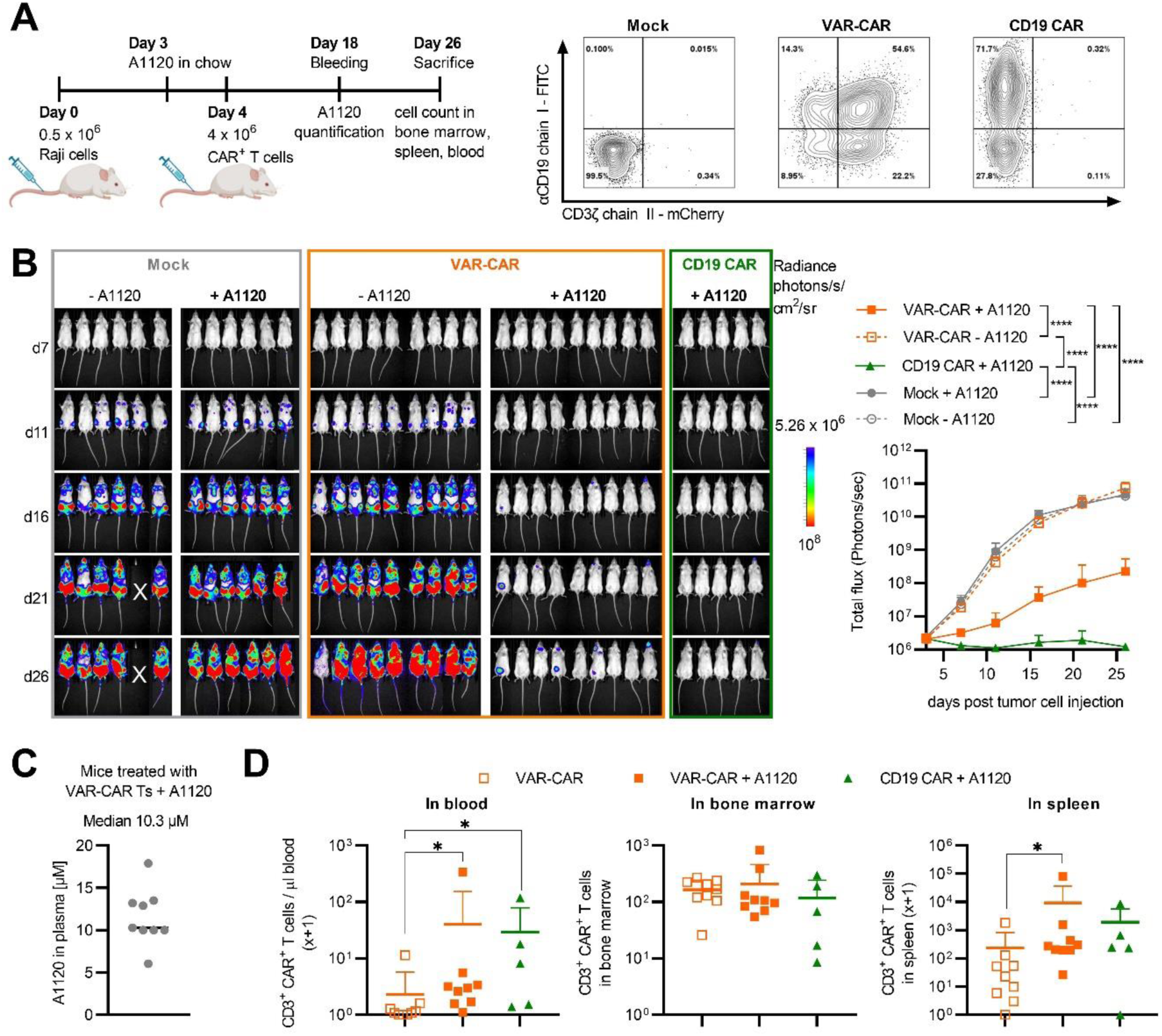
Regulation of VAR-CAR function by A1120 *in vivo*. **(A)** Schematic of the *in vivo* experiment and CAR expression in the cells injected into mice. **(B)** Pictures (left) and BLI curves (right) of the *in vivo* experiment. **** < 0.0001; 2-way Anova with Turkey’s test. Comparisons that did not reach statistical significance are not indicated. **(C)** Mass spectrometric analysis of A1120 concentration in the plasma of mice treated with A1120 and VAR-CAR T cells. **(D)** CD3^+^ CAR^+^ T cell counts per µL blood (left), in bone marrow (middle; total number in both femurs), and in the spleen (right; total number in the entire organ). For the blood and spleen, values were plotted as “x+1” to enable their depiction on a log scale. * < 0.05; ** < 0.01; *** < 0.001. Kruskal-Wallis test with Dunn’s correction. Comparisons that did not reach statistical significance are not indicated.

Analysis of CAR T cell numbers in blood, bone marrow and spleen at the end of the experiment revealed a trend towards higher VAR-CAR T cell numbers in the blood and spleen of A1120-treated mice compared to those that did not receive A1120 (Fig. 5D, Extended Data Fig. 6B). Phenotypic characterization of CAR T cells showed no major differences between treatment groups, both in terms of T cell differentiation and expression of exhaustion markers (Extended Data Fig. 6C and 6D).

Taken together, these results demonstrate that the anti-tumor potency of VAR-CAR T cells can be tightly controlled *in vivo* via oral administration of A1120.

### Integration of V35 and RF2 into a synthetic transcription factor to regulate gene expression

To further investigate the applicability of the V35-A1120-RF2 switch, we integrated the switch into a synthetic transcription factor to regulate gene expression. For this, we fused V35 to the N1 zinc finger DNA binding domain^35^ (DBD) and RF2 to the p65 activation domain (AD) in both orientations (i.e. N-terminal and C-terminal fusions, Fig. 6A). In addition, we included in the reverse strand a cassette for the expression of an anti-CD20 CAR driven by a minimum promoter preceded by ten repeats of N1 zinc finger binding sites.

**Figure 6.**
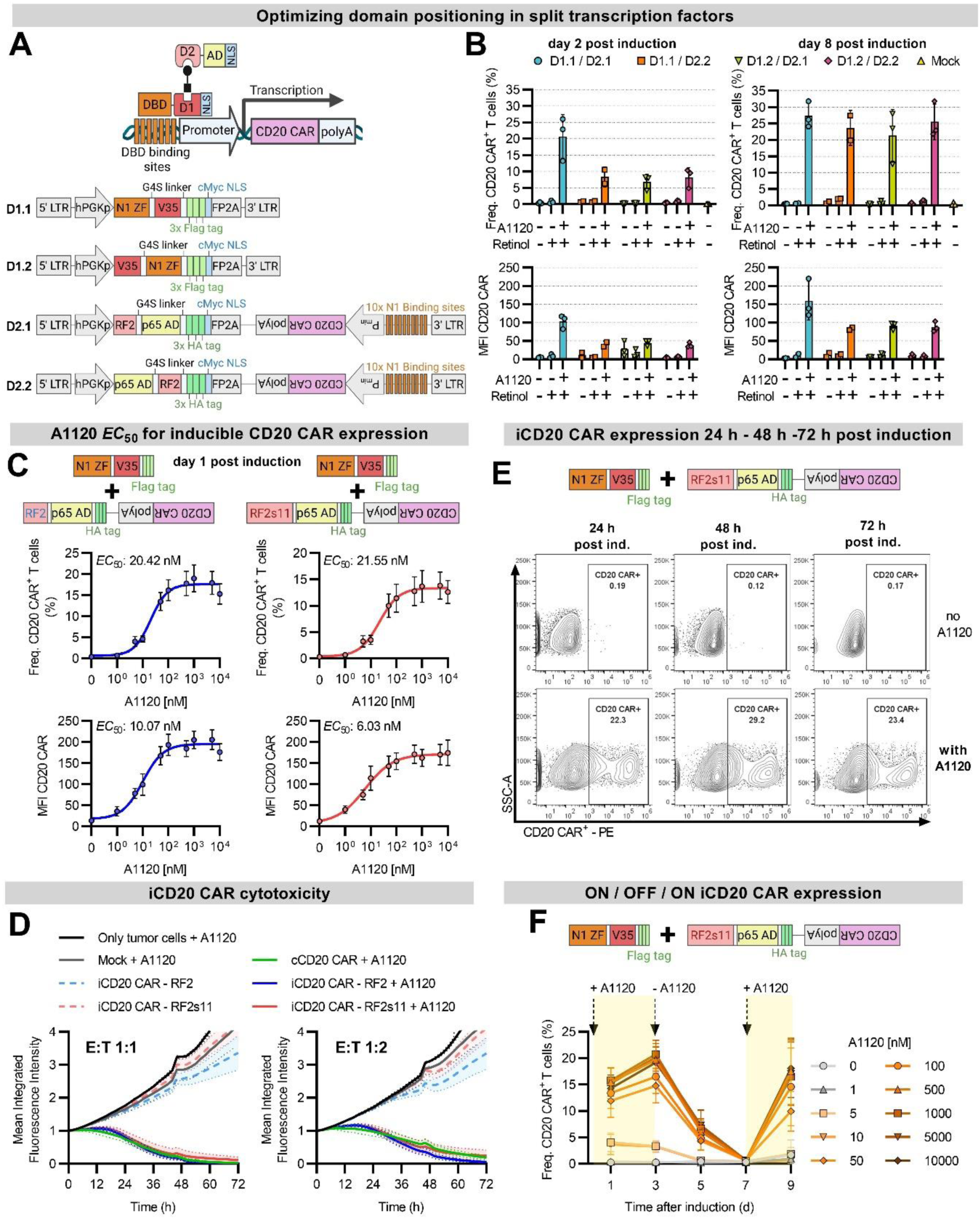
Regulation of gene expression by A1120 *in vitro*. **(A)** Schematic of the split transcription factor system and the different constructs. AD: activation domain. DBD: DNA binding domain. NLS: Nuclear localization domain. FP2A: Furin + P2A sites. D1 and D2: Dimerization domain 1 and 2. **(B)** Frequency of CD20 CAR positive cells (top) and CAR MFI (bottom) were measured two days (left) or eight days (right) after A1120 induction of CAR expression. A1120 (50 µM) and retinol (2 µM) were added as indicated (mean ± SD, n=3, 3 different T cell donors). **(C)** CD20 CAR expression one day after induction by increasing concentrations of A1120. The optimal domain positioning was used, combining either RF2 or RF2s11 with V35. Both the frequency of CD20 CAR^+^ T cells (top) and the CAR MFI (bottom) are shown (n=7, 7 different T cell donors). **(D)** CD20^+^ GFP^+^ 526-Mel target cells were co-cultured with the different CAR constructs at E:T 1:1 (left) or 1:2 (right) and target cell lysis was analyzed by live-cell imaging. The mean integrated fluorescence intensity was normalized to time = 0 h. 100 nM of A1120 were added to the indicated conditions at the start of the co-culture. (n=7, 7 different T cell donors). **(E)** Expression of an iCD20 CAR with a transcription factor containing RF2s11 measured 24, 48, and 72 hours after induction by 5 µM of A1120 or without the drug. **(F)** Frequency of CD20 CAR^+^ T cells upon addition and removal of different A1120 concentrations at the indicated time points (mean ± SD, n=4, 4 different T cell donors).

In cells containing both components of the split transcription factor and the CD20 CAR gene, the fusion protein of V35 and the N1 zinc finger DBD binds to the N1 zinc finger binding sites upstream of the promoter. However, expression of the CD20 CAR is only induced upon A1120-administration, which enables the p65 AD-RF2 construct to bind V35 that is fused to the DBD, thereby assembling a functional transcription factor and driving CAR expression (Fig. 6A).

In an initial experiment, we tested all combinations of the two split transcription factor components in primary human T cells, yielding the highest levels of inducible CD20 CAR expression with D1.1/D2.1 (Fig. 6B). Notably, the increased performance of D1.1/D2.1 was due to functional superiority instead of elevated expression levels of the subunits (Extended Data Fig. 7A and 7B). Moreover, in line with the yeast display data (Fig. 2D), we did not observe any activation of the switchable transcription factors upon addition of the natural hRBP4-ligand retinol or in the absence of A1120 (Fig. 6B).

Based on its potent A1120-induced CD20 CAR expression, we chose the D1.1/D2.1-based split transcription factor and termed this system “inducible CD20 CAR” (iCD20 CAR). We generated iCD20 CAR systems with D2.1 containing either RF2 or its stabilized variant RF2s11 and assessed their A1120-sensitivity in primary human T cells. Both versions were analyzed on day 1 and 3 post induction and with respect to the percentage of CAR^+^ cells and MFI, yielding highly consistent *EC*_50_ values in the range of ∼5-30 nM (Fig. 6C and Extended Data Fig. 7C). Next, we evaluated the anti-tumor efficacy of RF2- and RF2s11-based iCD20 CAR T cells in co-cultures with CD20^+^ GFP^+^ 526-Mel target cells. Both iCD20 CAR T cell versions showed potent anti-tumor activity in the presence of A1120 that was similar to that of CAR T cells expressing the same CD20 CAR constitutively via the EF1α promoter (cCD20 CAR, Fig. 6D and Extended Data Fig. 7D for individual curves of all 7 donors). In contrast, in the absence of A1120, RF2s11-based iCD20 CARs were completely inactive and comparable to Mock T cells, whereas their RF2-based counterparts showed some level of background killing. Based on this leakiness, which was only observed with RF2, but not with its stabilized version RF2s11, we chose the RF2s11-containing iCD20 CAR as the final system for further experiments.

Interestingly, high CD20 CAR expression levels are reached already within 24 h post A1120 administration and only slightly further increased after 48 or 72 h (Fig. 6E and 6F). Moreover, upon withdrawal of A1120 after an initial induction phase, CD20 CAR expression dropped to baseline levels within 4 days but could be efficiently reinduced (Fig. 6F).

Summing up, we established a split transcription factor system, which enables A1120-dependent CAR expression in human T cells. Once CAR expression is induced by the drug, these iCD20 CAR T cells efficiently eliminate tumor cells, while showing absolutely no background activation in the absence of A1120, demonstrating the tight control of this system.

### Drug-induced CAR expression and anti-tumor potency of iCD20 CAR T cells *in vivo*

Finally, we tested whether A1120-induced iCD20 CAR expression could be controlled *in vivo* via oral drug administration. In addition to their drug-dependency, we also assessed the anti-tumor potency of iCD20 CAR T cells in a stress test *in vivo* model, in which only 0.5×10^6^ CAR^+^ T cells were administered 5 days after Raji tumor cell injection (Fig. 7A).

**Figure 7.**
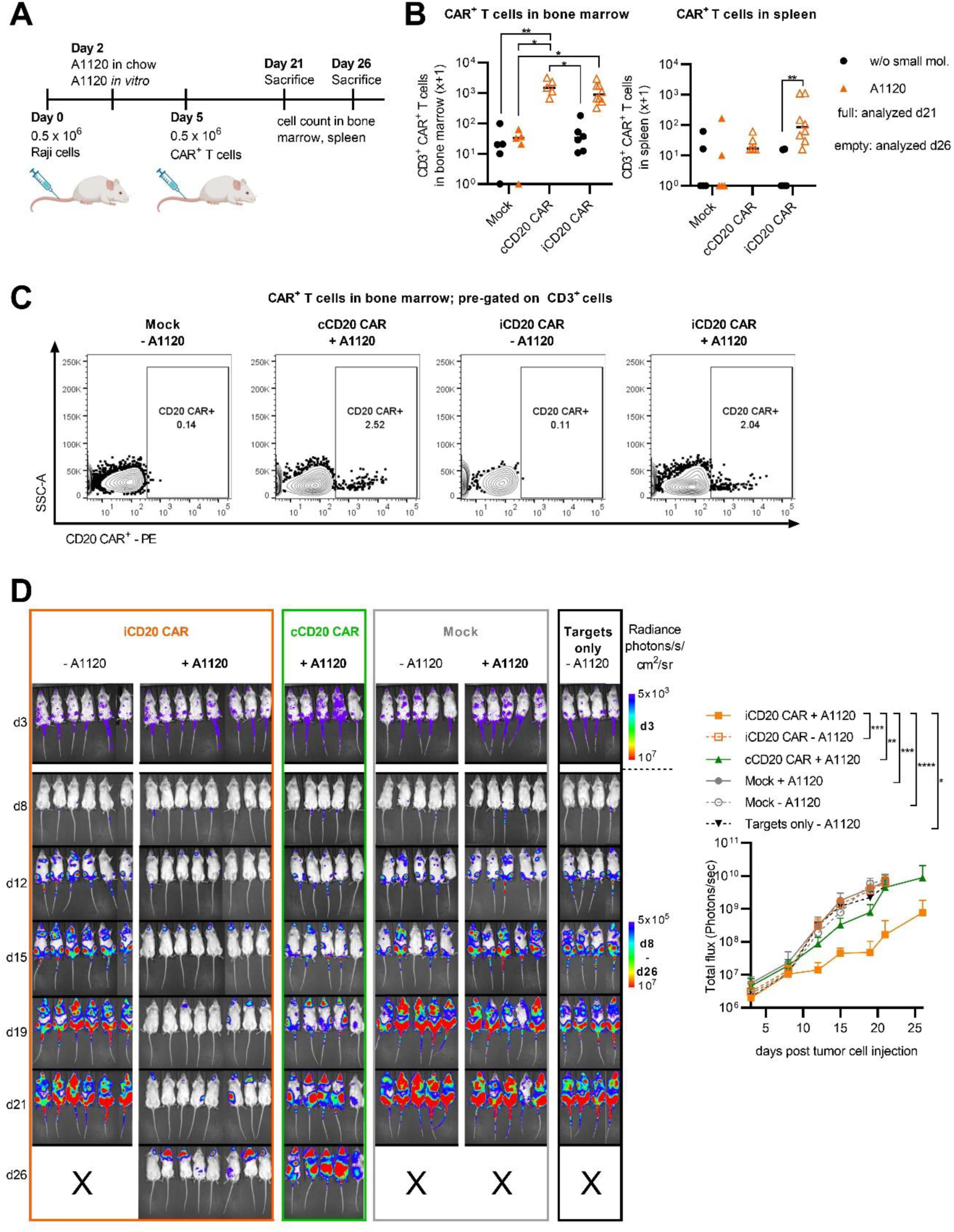
Regulation of CD20 CAR expression and anti-tumor activity by A1120 *in vivo*. **(A)** Schematic of the *in vivo* experiment. **(B)** CD3^+^ CAR^+^ T cell counts in bone marrow (left) and spleen (right). Values were plotted as x+1. * < 0.05; ** < 0.01. Kruskal-Wallis test with Dunn’s correction. Comparisons that did not reach statistical significance are not indicated. **(C)** Representative flow cytometry plots showing CAR expression in CD3^+^ T cells isolated from the bone marrow. **(D)** Images (left) and BLI curves (right) from the *in vivo* experiment. * < 0.05; ** < 0.01; *** < 0.001; **** < 0.0001. 2-way Anova with Turkey’s test. Comparisons that did not reach statistical significance are not indicated.

When we analyzed CAR T cell numbers at the end of the experiment, we detected similar numbers of CAR^+^ T cells in iCD20 CAR vs. cCD20 CAR T cell treated mice. Importantly, in line with our *in vitro* experiments, no CAR expression could be observed without administration of A1120, yielding background levels comparable to Mock T cells (bone marrow and spleen, Fig. 7B, 7C and Extended Data Fig. 8B). No major differences were found with respect to total T cell numbers (Extended Data Fig. 8B).

The A1120-induced iCD20 CAR expression also translated to drug-dependent anti-tumor activity. That is, mice receiving iCD20 CAR T cells without the drug showed uncontrolled tumor growth comparable with Mock T cells, whereas A1120 administration resulted in potent tumor elimination. Remarkably, in this *in vivo* stress test model, iCD20 CAR T cells showed even higher anti-tumor efficacy than the cCD20 CAR T cells (Fig. 7D).

Together, these data demonstrate that our A1120-responsive split transcription factor system can tightly regulate the expression of a CAR *in vivo*, yielding potent tumor control upon oral administration of A1120.

## Discussion

In this study, we engineered a drug-responsive switch that combines multiple critical advantages for applications in cellular therapies: (i) protein components based on human sequences (hRBP4 and Fn3 domain), to reduce the risk of immunogenicity; (ii) the orally available small molecule drug A1120^26–29,36^, facilitating convenient administration without the need for any medical infrastructure or equipment (in contrast to i.v. injections); (iii) a good safety profile of the drug with no toxicities observed even at high dose and after long-term application in mice for 6 months^29^; (iv) intracellular and extracellular functionality, thus expanding the range of potential applications; (v) stable and well-expressed protein components of small size and (vi) tight control by the drug.

We demonstrated the modularity and versatility of our newly developed heterodimeric switch by testing it in two different applications, including membrane-bound split CARs, as well as cytoplasmic/nuclear split transcription factors regulating expression of a gene of interest, such as a CAR. When tested *in vivo*, both systems were tightly controlled, showing no background activation in the absence of A1120. Importantly, oral A1120 administration induced potent tumor control by either assembling a split CAR (VAR-CAR) or by inducing expression of the CAR (iCD20 CAR). iCD20 CAR T cells were even more potent compared with those expressing the same CAR constitutively (cCD20 CAR). This superior anti-tumor function is unlikely a result of elevated CAR T cell levels in the iCD20 CAR group, since these numbers were comparable between the cCD20 and iCD20 expressing groups (Fig. 7B, 7C and Extended Data Fig. 8B). Instead, it might be explained by the slightly increased CAR expression densities found on iCD20 CAR T cells (Extended Data Fig. 8A) or by reduced T cell exhaustion due to the lack of CAR expression during *in vitro* expansion. In fact, similar effects have been observed with SynNotch CAR T cells, where the CAR is also absent during *in vitro* manufacturing^37,38^.

To sum up, we engineered a fully human switch that is efficiently regulated with a safe and orally available small molecule drug. We demonstrated its functionality in two highly relevant applications, i.e. control of (i) CAR function and (ii) transcriptional activity. These switchable systems may address multiple limitations in the CAR T cell field, including the clinical management of toxicities, control of CAR T cell expansion – in particular with next generation CAR T cells engineered for enhanced persistence and proliferation, reduction of CAR T cell exhaustion and/or inactivation of CAR T cells after successful therapy, thereby reversing e.g. B cell aplasia in the case of CD19-directed CAR Ts^39^. Of note, the switchable transcription factors presented in this study can be used to regulate the expression of virtually any gene of interest, which opens up numerous potential applications for cell therapies. Therefore, we anticipate that the A1120-responsive switch developed in this study will be a critical step towards the development of next-generation cellular therapeutics with improved safety and efficacy.

## Acknowledgements

This work was supported by the Austrian Federal Ministry of Economy and Tourism, the National Foundation for Research, Technology and Development, the Christian Doppler Research Association (Christian Doppler Laboratory for Next Generation CAR T Cells) and by private donations to the St. Anna Children’s Cancer Research Institute (Vienna, Austria). This research was also funded in part by the Rete ACC - Sviluppo successivo per il progetto CAR-T rete Oncologica: ponte alla traslazione clinica (RCR-2023-23684268) and the Austrian Science Fund (FWF) [W1224–Doctoral Program on Biomolecular Technology of Proteins–BioToP; EFP 45, Devising Advanced TCR-T cells to eradicate OsteoSarcoma (DART2OS) and FWF grant 10.55776/PAT8789924]. For open access purposes, a CC BY public copyright license has been applied to any author who accepted manuscript version arising from this submission. E.S. is a recipient of DOC Fellowships of the Austrian Academy of Sciences at the St. Anna Childreńs Cancer Research Institute (#26323). Research at the IMP is supported by Boehringer Ingelheim and the Austrian Research Promotion Agency (headquarter grant FFG-852936). M.S. is a member of the Boehringer Ingelheim Discovery Research global post-doc program. We further acknowledge the CCRI FACS Core Unit for providing cell sorting and analysis services, the Connective Base GmbH, BOKU Core Facility Biomolecular & Cellular Analysis and BOKU Core Facility Mass Spectrometry for providing flow cytometers, MS equipment, and supporting the project. All schematic figures were created with Biorender.com.

## Data availability

Source data will be provided with this paper and deposited in a publicly available repository before final publication.

## Author contributions

E.S., M.W.T. and M.L. conceived the study. E.S., A.M-L., G.D., D.E., K.M., F.E., H.B., S.B., T.M., M.S., performed experiments and acquired data. E.S., A.M-L., G.D., D.E., K.M., F.E., H.B., S.B., T.M., D.M., A.U., M.S., J.M., J.Z., A.R., B.E., C.U.Z., M.W.T., M.L. designed experiments. E.S., A.M-L., G.D., D.E., K.M., B.E., C.U.Z., M.W.T., M.L. analyzed data and interpreted results. D.M. and A.U. developed the method for mass spectrometric quantification of A1120 in mouse plasma, performed the measurements and data analysis. B.E., M.W.T. and M.L. supervised the work. E.S. and M.W.T. wrote the manuscript. All authors edited and approved the manuscript.

---

**Extended Data Fig. 1:**
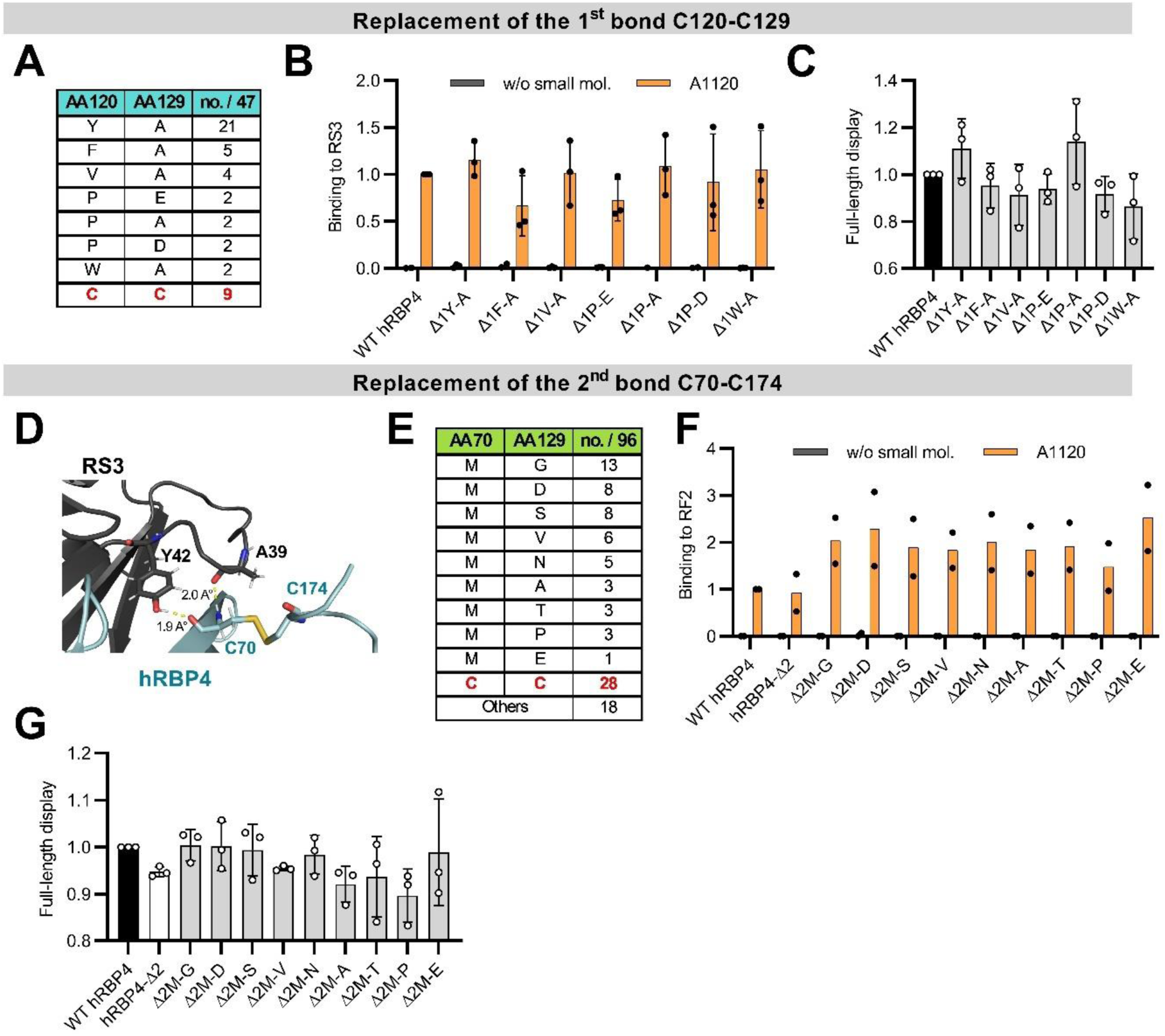
Replacement of the first two disulfide bonds. **(A)** Mutants enriched after directed evolution of hRBP4-**Δ1NNK** are shown. The frequency among the 47 sequenced clones is given in the third column as the number of clones found (no.) out of 47 sequenced.2-way Anova with Turkey’s test. **(B)** Binding of the hRBP4-Δ1NNK variants to 300 nM RS3 +/− 5 µM A1120. gMFI (geometric mean fluorescence intensity) values were normalized to the binding of WT hRBP4 in the presence of A1120 (mean ± SD, n=3). **(C)** Full-length expression of hRBP4-Δ1NNK variants on the surface of yeast cells as measured via c-myc tag detection. Expression levels (gMFI) were normalized to WT (mean ± SD, n=3). **(D)** Close-up view of the crystal structure of hRBP4 loaded with A1120 and bound to RS3 (PDB: 6QBA^13^. Potential hydrogen bonds between C70 of hRBP4 and Y42 and A39 of RS3 (1.9 A° and 2.0 A° respectively) are indicated in yellow dashed lines. **E)** Mutants enriched through directed evolution. The frequency among the sequenced clones is given in the third column as the number of clones found (no.) out of 96 sequenced. “Others” refers to non-enriched variants that are either truncated or contain additional point mutations outside positions 70 and 174. **F)** Binding of hRBP4-Δ2 NNK variants to 1 µM RF2 binder +/− 5 µM A1120 (n=2). gMFI values were normalized to WT hRBP4 in the presence of A1120. **G)** Full-length expression of hRBP4-Δ2 NNK variants on the surface of yeast cells. Expression levels (gMFI) were normalized to WT (mean ± SD, n=3).

**Extended Data Fig. 2:**
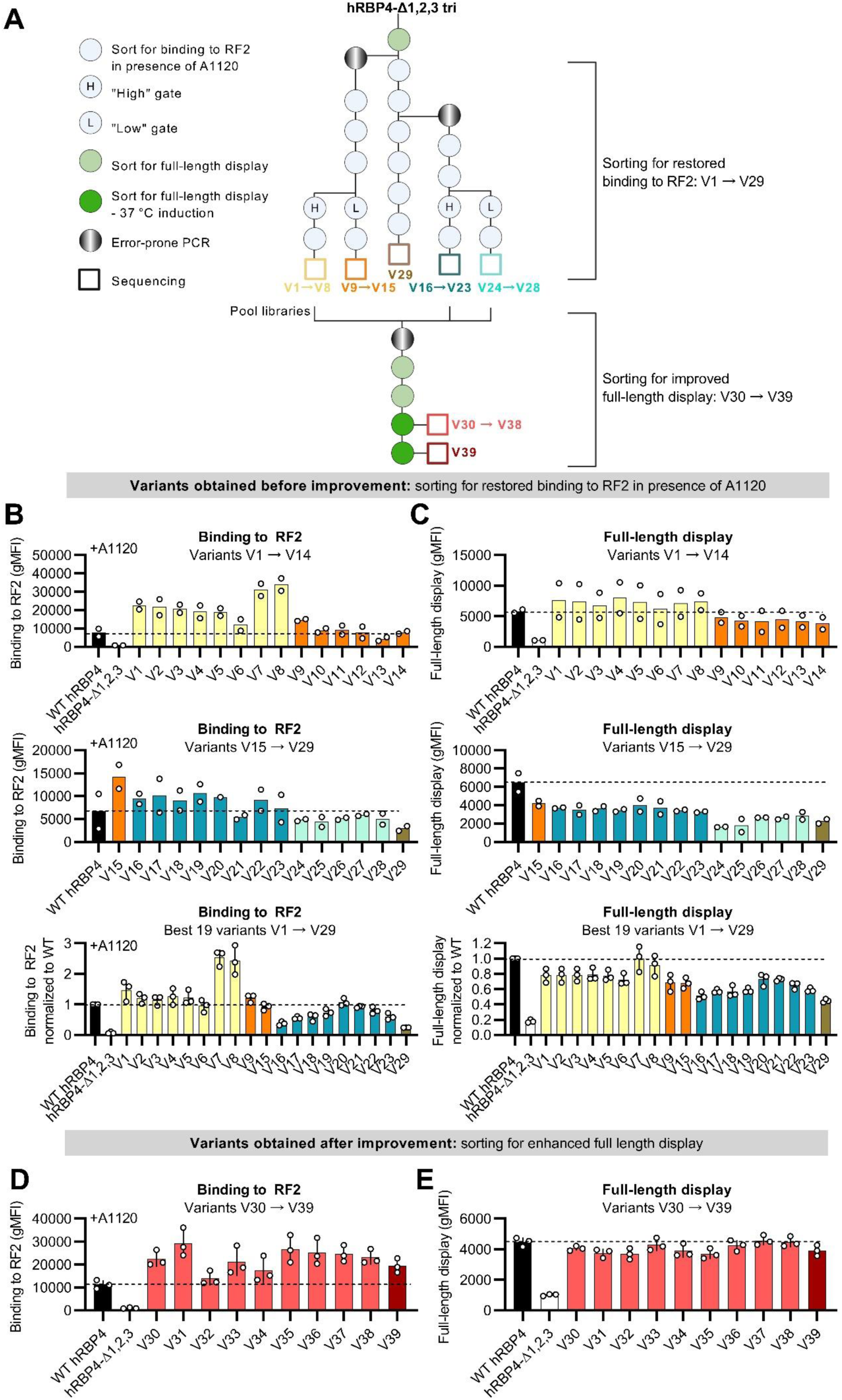
Engineering of the third disulfide bond and high throughput screening of obtained variants. **A)** Schematic for the engineering of the last library ‘hRBP4-Δ1,2,3 tri’. **(B to E)** Preliminary screening of the dfRBP4 variants obtained before (A and B) or after (C and D) improvement. **(B and D)** Binding of the disulfide-free hRBP4 (dfRBP4) variants to 1 µM RF2 in the presence of 5 µM A1120 (n=2 or mean ± SD and n=3). **(C and E)** Full-length expression of the dfRBP4 variants on the surface of yeast cells (n=2 or mean ± SD and n=3).

**Extended Data Fig. 3:**
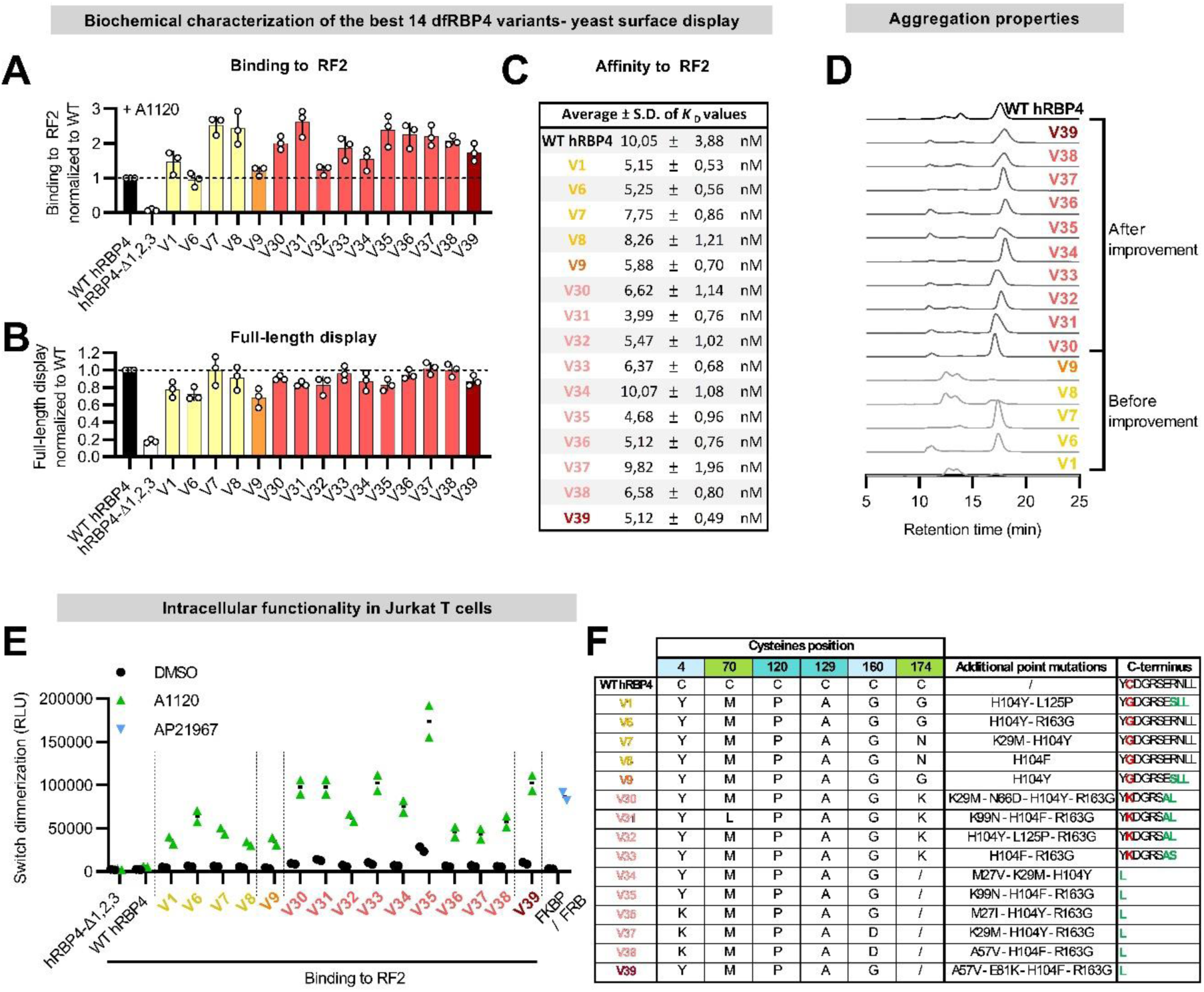
Characterization of the 15 best dfRBP4 variants on yeast. **(A)** Binding of the dfRBP4 variants to 1 µM soluble RF2 in the presence of 5 µM A1120, normalized to the binding of WT hRBP4 (mean ± SD, n= 3). **(B)** Full-length expression of the disulfide-free variants on the surface of yeast cells. Expression levels were normalized to WT (mean ± SD, n=3). **C)** Affinity of the dfRBP4 variants to RF2. *K*_D_ values were measured by titrating soluble RF2 onto yeast-displayed dfBRP4 variants in the presence of 5 µM A1120 (mean ± SD, n=3). **D)** Size exclusion chromatography profile of dRBP4 variants expressed solubly in the cytoplasm of *E.coli*. One representative measurement of three independent experiments is shown. **E)** Analysis of A1120-induced dimerization of cytoplasmic dfRBP4 variants and RF2 using the NanoBiT® assay. A1120 (5 µM) was added as indicated. Binding of FKBP to FRB in the presence of rapalog (AP21967, 0.5 µM) was used as a benchmark. RLU: relative light units. **F)** Sequences of the best 15 dfRBP4 variants compared to the WT hRBP4, which is shown as reference in the first line. The colors in the first column represent the sub-libraries from which the variants were derived.

**Extended Data Fig. 4:**
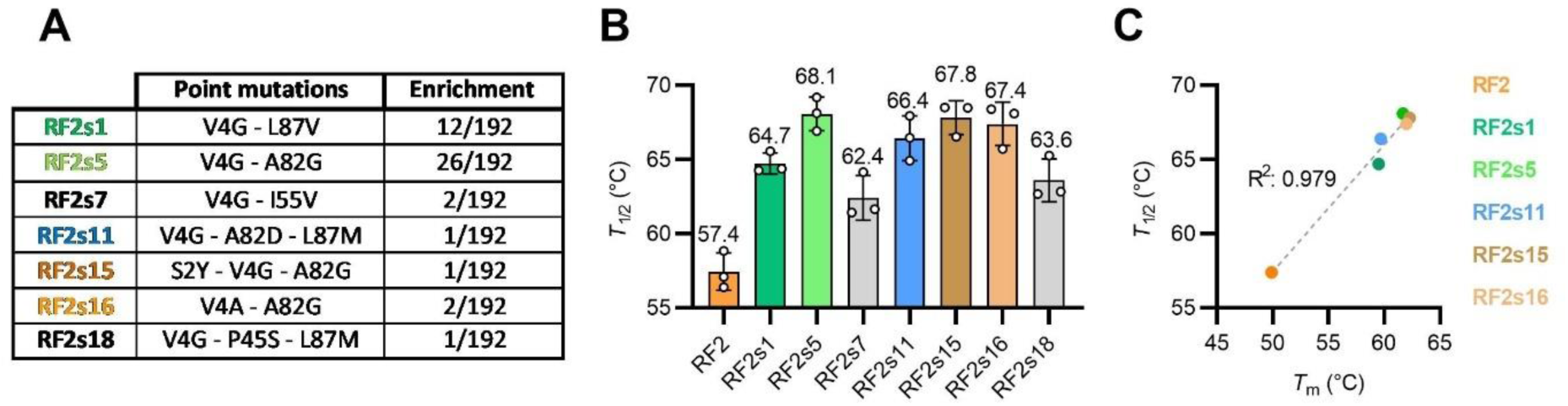
Characterization of the 7 best stabilized RF2 variants. **(A)** Point mutations of the 7 best stabilized RF2 variants compared with non-stabilized RF2. **(B)** *T*_1/2_ values of stabilized RF2 mutants. Binding to dfRBP4 V35 was used to monitor RF2 unfolding in heat-exposed yeast cells (stained after heat shock with 20 nM dfRBP4 V35 and 5 µM A1120) (mean ± SD, n=3). **(C)** Correlation between *T*_1/2_ and *T*_m_ values shown in Figure 3 D for the different RF2 stabilized variants.

**Extended Data Fig. 5:**
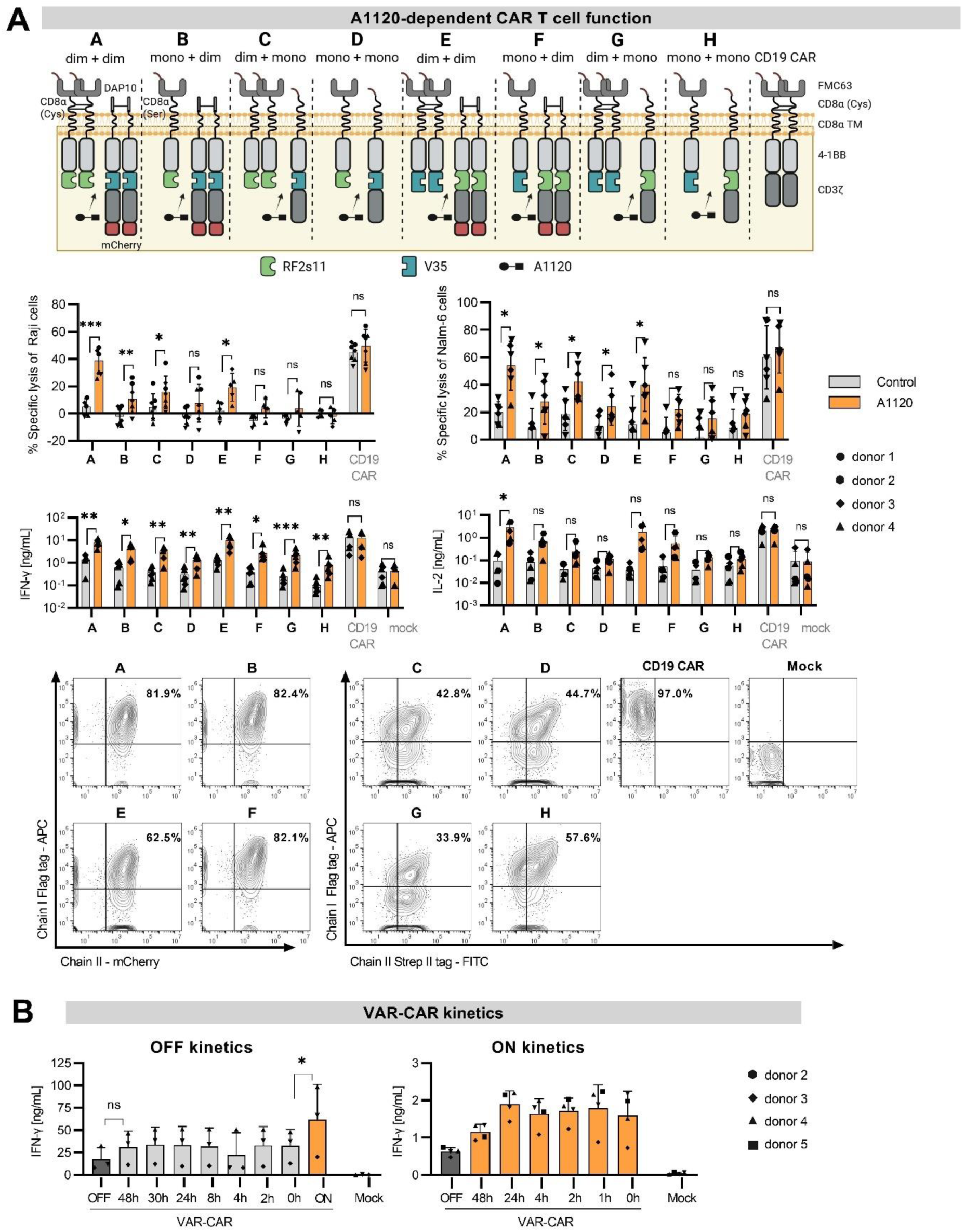
Regulation of split CAR function by A1120 *in vitro*. **(A)** Comparison of different split CAR designs. A schematic of the different split CAR architectures tested is shown at the top, lysis of Raji (E:T 2:1, 24 h, mean ± SD, n=5-7) and Nalm-6 cells (E:T 2:1, 6 h, mean ± SD, n=6), as well as IFN-γ (mean ± SD, n=6) and IL-2 (mean ± SD, n=5) secretion after co-culture with Raji cells, are shown in the middle. Expression of the different CAR constructs is shown at the bottom. For the dimeric chain I (chain I dimer), the CD8α hinge with two cysteines was used. Mutation of these cysteines to serines resulted in monomeric chain I (chain I mono). The monomeric chain II (chain II mono) has an extracellular StrepII-tag fused to a CD8α hinge with cysteine-to-serine mutations. To dimerize chain II (chain II dimer), the CD8α hinge was exchanged with DAP10. mCherry was fused C-terminally after the CD3ζ domain for detectability of expression. **(B)** OFF (left) and ON (right) kinetics of VAR-CAR activation. For the OFF kinetics, VAR-CAR T cells were cultured with 5 µM A1120 (72 h before the co-culture) and subsequently washed to remove the drug at the indicated time points before co-culture with Raji cells without A1120 (E:T of 2:1, 24 h). The time points represent the duration of culture in the absence of A1120 before co-culture with Raji cells. VAR-CAR cells continuously exposed to A1120 were washed once and A1120 was added (ON) or not added (0 h) in the co-culture. Mock T cells and VAR-CAR cells never cultured with A1120 (OFF) served as negative controls. IFN-γ secretion was measured in the supernatants after 24 hours (n = 3, three different T cell donors). For the ON kinetics, 5 µM A1120 was added to VAR-CAR T cell cultures at the indicated time points prior to the co-culture with Raji target cells. At t = 0 h, a 4 h co-culture with Raji cells was initiated in the presence of A1120(E:T of 2:1). Mock T cells and VAR-CAR cells never cultured with A1120 (OFF) were used as negative controls. IFN-γ secretion was measured in the supernatants after 4 hours (n = 4, 4 different T cell donors). * < 0.05; ** < 0.01; *** < 0.001. Multiple paired t-test (lysis) or ratio t-test (cytokine) with Holm Sidak correction.

**Extended Data Fig. 6:**
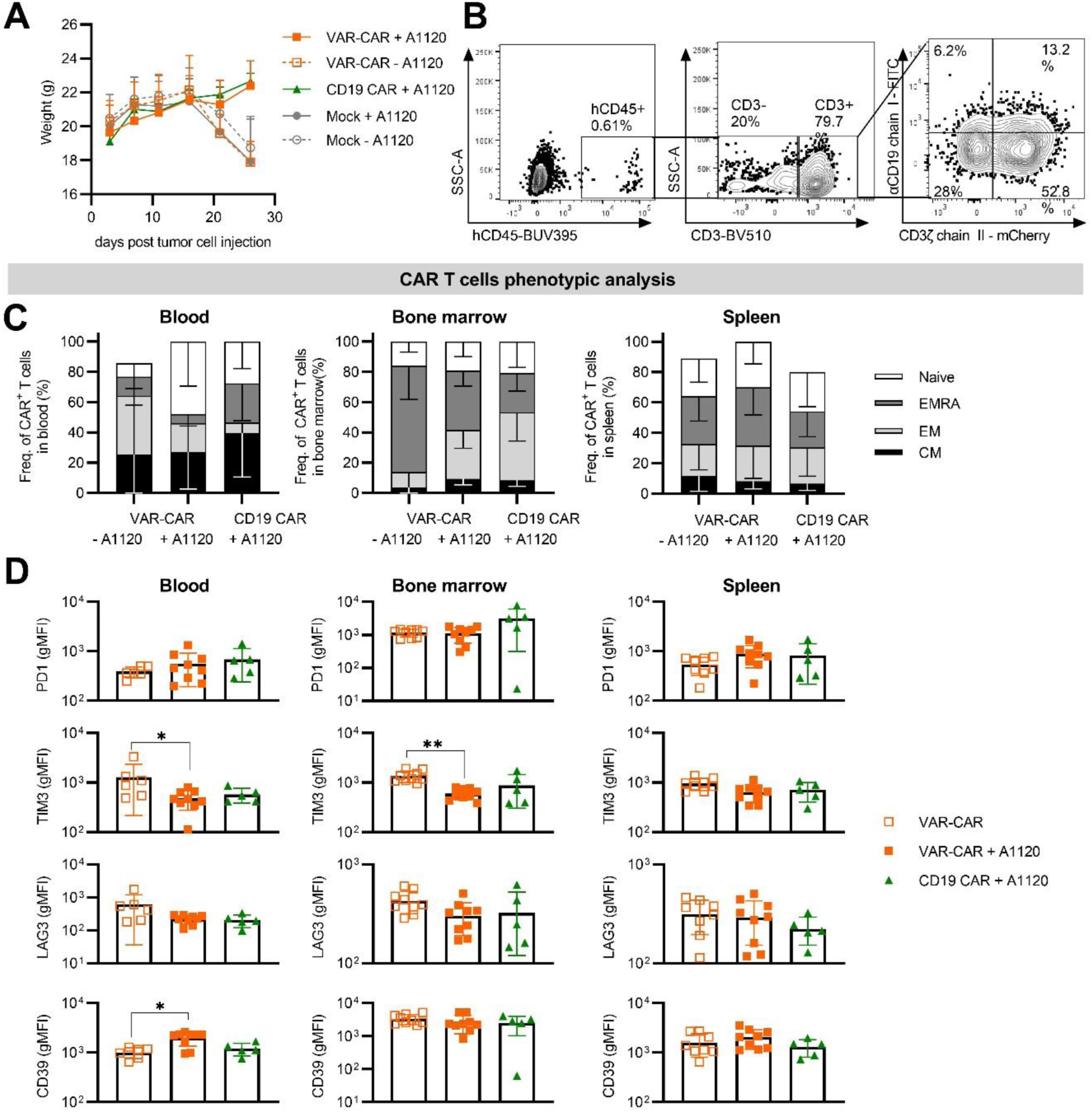
Regulation of split CAR function by A1120 *in vivo*. **(A)** Weight of the mice during the experiment. **(B)** Gating strategy for the cell counts and phenotypic analysis **(C)** CD3^+^ T cell counts per µL of blood (left), in the bone marrow (middle) and in the spleen (right), pre-gated on CD45. **(D)** and (E) Phenotypic analysis of the CAR+ cells in blood, bone marrow and spleen for the VAR-CAR and the CD19 CAR, with (D) the differentiation phenotype (CM: central memory; EM: effector memory; EMRA: effector memory with expression or RA) and (E) the expression of exhaustion markers PD1, TIM3, LAG3 and CD39. Phenotypic analysis of the * < 0.05; ** < 0.01; *** < 0.001. Kruskal-Wallis test with Dunn’s correction. Comparisons that did not reach statistical significance are not indicated.

**Extended Data Fig. 7:**
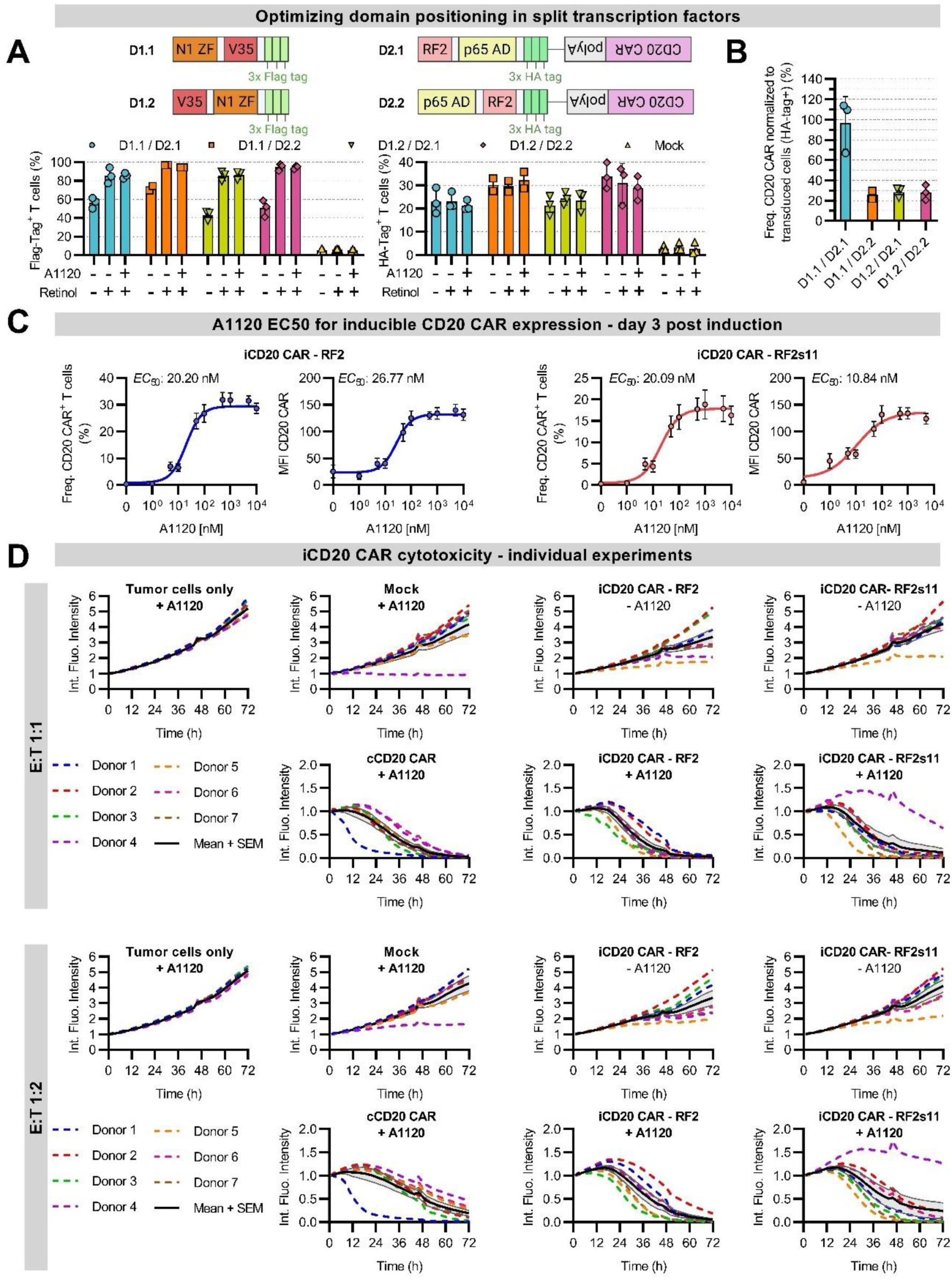
Regulation of CD20 CAR expression by A1120 *in vitro*. **(A)** Percentages of Flag (left) and HA (right) positive cells two days after A1120 induction of CAR expression. A1120 (50 µM) and retinol (2 µM) were added as indicated (mean ± SD, n=3, three different T cell donors). **(B)** Frequency of CD20 CAR positive cells normalized to HA-positive cells for the different domain combinations two days after A1120 induction. **(C)** CD20 CAR expression measured three days after induction by increasing concentrations of A1120. The D1.1/D2.1 domain positioning was used with either RF2 (blue) or RF2s11 (red) in combination with V35. Both the frequency of CD20 CAR^+^ T cells and the CAR MFI are shown (n=7, 7 different T cell donors). **(D)** Individual curves of the integrated fluorescence intensity from the experiment shown in Figure 6D for E:T ratios of 1:1 (top) and 1:2 (bottom).

**Extended Data Fig. 8:**
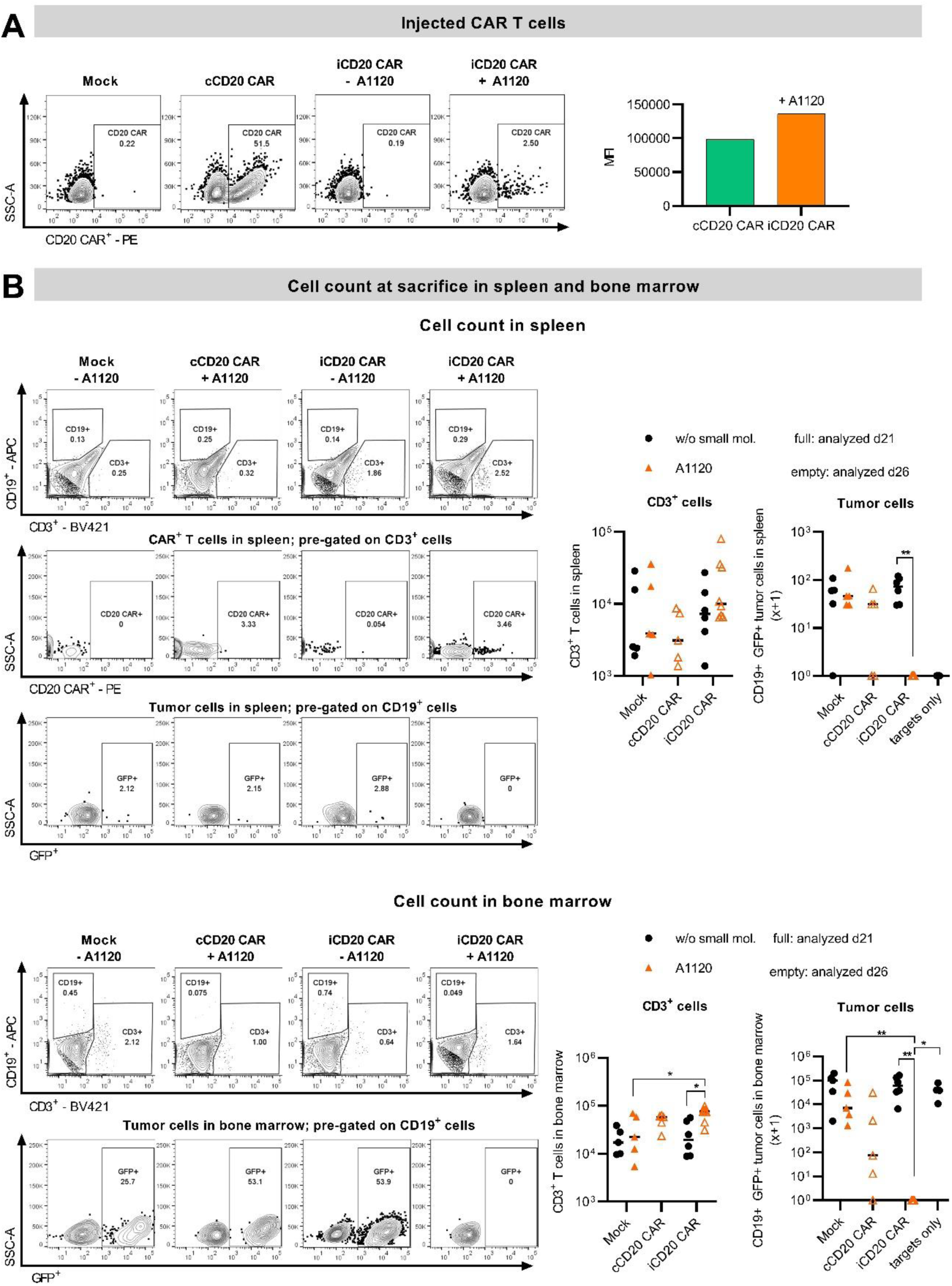
Regulation of CD20 CAR expression and anti-tumor potency by A1120 *in vivo*. **(A)** CAR expression frequency (left) and MFI (right, gated on CAR^+^) of the T cells injected into mice. **(B)** Gating strategy and representative flow cytometry plots, as well as CD3^+^ T cell and tumor cell counts in the spleen (top) and bone marrow (bottom). CAR^+^ T cell counts for both organs and representative flow cytometry plots for the bone marrow are shown in Figure 7. * < 0.05; ** < 0.01; Kruskal-Wallis test with Dunn’s correction. Comparisons that did not reach statistical significance are not indicated.

## Materials and Methods

### Recombinant expression of proteins

The different RBP4 variants were expressed recombinantly as soluble proteins fused to an N-terminal 6x His-tag using the pET21+ vector. The stabilized RF2 mutants were fused to an N-terminal 6x His-tag and a SUMO protein domain using the pE_SUMO vector. The constructs were transformed into *Escherichia coli* Rosetta cells for disulfide-free RBP4 and RF2 stabilized variants or Origami cells for wild-type RBP4. After overnight incubation in lysogeny broth (LB) medium supplemented with ampicillin and chloramphenicol (100 µg/mL) (chloramphenicol was not added for the Origami cells) at 37 °C, 180 rpm, cultures were set to an optical density at 600 nm (OD_600_) of 0.2 and further incubated at 37 °C. Protein expression was induced by addition of 1 mM of isopropyl β-D-1-thiogalactopyranoside (IPTG) when the OD_600_ reached 1, followed by overnight incubation at 20 °C. Cells were harvested the next day by centrifugation (5000 g, 20 min, 4 °C). The pellet was resuspended in sonication buffer (50 mM phosphate buffer pH 7.5 with 300 mM NaCl, 3 % glycerol, 1 % Triton X-100) and the cells were lysed by sonication (2 min, pulse: 1,0 – 1,0 seconds, 90 % amplitude) on ice. The cell debris were removed by a centrifugation step (20 000 g, 30 min at 4 °C) and the supernatant containing the soluble proteins was supplemented with 10 mM imidazole and filtered through a 0.45 µM filter (Duropore).

6x His-tagged proteins were purified by gravity flow with a TALON metal affinity resin (Takara Bio). The resin was equilibrated (equilibration buffer: 50 mM sodium phosphate, 300 mM NaCl, pH 8) and the supernatants were loaded twice. The same buffer containing increasing concentrations of imidazole was added to the column for multiple washing steps (5 mM and 15 mM imidazole) and elution of the proteins (250 mM imidazole). For large-scale expression, 6x His-tagged proteins were purified by immobilized metal affinity chromatography using a HisTrap FF column connected to a Bio-Rad purifier system. After loading of the supernatant and washing steps (washing buffer: 50 mM sodium phosphate, 500 mM NaCl, 5 mM imidazole, pH 7.4), elution of the pure proteins was carried out by applying a linear imidazole gradient (from 5 % to 100 % of an elution buffer containing 500 mM imidazole). Absorbance at 280 nM was detected and the fractions containing a protein of the right molecular weight (as analyzed via SDS-PAGE) were pooled. Buffer exchange to PBS was performed overnight at 4 °C via dialysis using a 10 kDa SnakeSkin dialysis tubing (Thermo Scientific). Purified proteins were concentrated via subsequent centrifugation steps in Amicon Ultra-15 10 K centrifugal filters (Merck Millipore).

Fractions of proteins were labelled with biotin using the EZ-Link Sulfo-NHS-LC-LC-Biotin kit (Thermo Scientific) according to the manufacturer’s instructions.

### Size Exclusion Chromatography (SEC)

Protein aggregates or unbound biotin molecules were removed via preparative size exclusion chromatography. After a centrifugation step (16 000 rpm for 3 min) the supernatant was loaded onto a HiLoad Superdex 75 pg column connected to a Bio-Rad NG-C chromatography system. PBS was used as running buffer and the fractions containing monomeric proteins based on absorbance at 280 nm were pooled and concentrated as mentioned above.

### High pressure liquid chromatography

Non-SEC purified proteins were centrifuged at 3000 g for 3 min to remove large aggregates and diluted to 1 mg/mL in PBS with 200 mM NaCl. The diluted samples were filtered through Ultrafree 0.1 µm centrifugal filter units by centrifuging them at 20,000 g for 5 min. The samples were then loaded to a Superdex 200 10/300 GL column connected to a HPLC Prominence LC20 system with a flow rate of 0.75 mL/min at 25 °C and a UV/VIS Photodiode Array Detector enabled detection of proteins via their absorbance at 280 nm.

### Differential scanning calorimetry

50 µM of the respective SEC purified proteins in PBS were heated up from 20 °C to 110 °C with a rate of 1 °C / min. Buffer runs without or with 1 µM A1120 (#A3111 Sigma, dissolved to 10 mg/mL in DMSO) for dfRBP4 and RF2 variants respectively were used for baseline subtraction. After normalization for protein concentration, transitions were fitted with a non-two state thermal unfolding model software to calculate melting temperatures (*T*_m_) using the MicroCal PEAQ-DSC.

### Surface plasmon resonance

20 µg/mL RF2s11-SUMO in 10 mM NaAC buffer pH 4.5 were covalently immobilized on a CM5 sensor chip (GE healthcare) via amine coupling according to the supplier’s instructions with a flow rate of 30 µL / min to a density of 2,000 response units (RU). Multi-cycle kinetic experiments were performed using increasing concentrations of dfRBP4 V35 in the presence or absence of 1 µM A1120 in the running buffer. When assessing binding in the presence of A1120, dfRBP4 V35 (3.2 nM to 125 nM) was pre-incubated with 5 µM A1120 for 1 h at 4 °C. To assess binding without the drug, dfRBP4 V35 was used at higher concentrations, ranging from 51.2 nM to 12.5 µM. The contact time and dissociation time were always 60 s and the flow rate 30 μL/min. 3 M MgCl_2_ was used to regenerate between cycles. The baseline bulk flow response was subtracted from the equilibrium RU and the resulting values were plotted against analyte concentrations. The data were then fitted to a 1:1 binding model to determine *K*_D_ values.

### Yeast surface display library preparation and screenings

The hRBP4-Δ1 NNK and hRBP4-Δ2 NNK gene libraries derived from hRBP4 WT with C120-C129 and C170-C174 mutated to NNK respectively were ordered from Twist.

For the hRBP4-Δ3 tri library an equimolar mix of 3 gene fragments containing the C70M, C120P and either C129A, C129E or C129D substitutions was amplified with a mix of 500 nM flanking primers (Fwd_1: 5’-GGCTCTGGTGGAGGCGGTAGCGGAGGCGGAGGGTCGGCTAGC and Rev_1: 5’-CTATTACAAGTCCTCTTCAGAAATAAGCTTTTGTTCGGATCC) and 20 nM of tri-nucleotide primers (Ella Biotech) (Fwd_2: 5’ - GCGGAGGCGGAGGGTCGGCTAGCGAGCGCGAC**X01**CGAGTGAGCAGCTTCCG AGTCA AG and Rev_2: 5’ - GTAACCGTTGTGGACGATCAGCCTGTACTGCCTGGCCAG**Z01**CAGCTCCTCCTG CCGCT GCCTTAC. X01 and Z01 encode for all amino acids except cysteines). To generate the RF2 library, wild-type RF2 DNA was subjected to an error-prone PCR (GeneMorph II Kit with 500 ng target DNA, 20 cycles).

Subsequent insert and vector (pCTCON3) preparation steps for all libraries as well as electroporation of the library in electrocompetent *S. cerevisiae* cells (strain EBY100, ATCC) were prepared according to Chen et al^40^. Library diversities were calculated by plating serial dilutions of the cells on selective SD-CAA plates after electroporation and counting transformants.

After electroporation, yeast libraries were cultivated to an OD_600_ of 1 in SD-CAA (20 g/L glucose, 6.7 g/L yeast nitrogen base, 5 g/L casamino acids, 11.85 g/L sodium citrate dihydrate and 7.4 g/L citric acid monohydrate) at 30 °C, 180rpm. Protein expression on the yeast surface was induced by switching the medium to SG-CAA (2 g/L glucose, 20 g/L galactose, 6.7 g/L yeast nitrogen base, 5 g/L casamino acids, 10.2 g/L disodium hydrogen phosphate and 4.82 g/L sodium phosphate monobasic) and incubating overnight at 20 °C. To select dfRBP4 variants binding to RF2 in the presence of A1120, the yeast library was incubated with 5 µM A1120 (30 min at 4 °C) before adding 1 µM soluble RF2 (1 h at 4 °C). A1120 (#A3111, Sigma) was dissolved to 10 mg/mL in DMSO and further diluted to the desired concentration in the proper medium. Biotinylated and non-biotinylated RF2-SUMO were used in alternative selection rounds to prevent selection of biotin binders. After a washing step with PBSA (PBS with 1 g/L BSA), the cells were stained for 20 min at 4 °C with either Penta-His-Alexa Fluor 647 (Qiagen) or streptavidin-Alexa Fluor 647 (Invitrogen) to measure binding to RF2, anti-HA-Alexa Fluor 488 (clone 16B12, Biolegend) to assess display or anti-c-myc AlexaFluor 488 (9E10, Thermofisher) to assess full-length display of the dfRBP4 variants. The anti-HA staining was always used to normalize binding or full-length expression signals to yeast cells displaying the protein. After a final washing step, the cells were sorted by fluorescence activated cell sorting (FACS), using a FACS Aria™ Fusion cell sorter or SONY Sorter SH800.

To enrich in dfRBP4 variants with improved thermostability, protein expression was induced at 37 °C instead of 20 °C. Cells were then incubated with both anti-c-myc AlexaFluor 647 and HA-Alexa Fluor 488 and for 20 min at 4 °C and sorted for improved full-length expression (higher c-myc signal).

To select thermostable RF2 variants, yeast cell displaying RF2 variants were subjected to a heat shock, i.e. they were first incubated on ice for 10 min, then on a thermocycler for 10 min (60 to 70°C depending on the sorting round) and once again on ice for 10 min. In parallel, 20 nM biotinylated or not biotinylated soluble dfRBP4 V35 was pre-incubated with 5 µM A1120 for 30 min at 4°C. After heat shock, the yeasts were centrifuged (2000 g, 3 min, 4°C), resuspended in buffer containing V35/A1120 and incubated for 1 hour at 4°C before proceeding with the usual staining. Since heat shocks above 48°C massively decrease cell viability, the cells cannot be directly cultivated after sorting. Here, the plasmid DNA pool in the sorted cells was isolated (Zymoprep yeast plasmid miniprep kit II, Zymo Research), further amplified with PCR and re-electroporated into new yeast cells with the linearized vector to generate a new library. Sorting rounds without heat shock were performed to enrich in living yeast cells.

After successful enrichments, the yeast library DNA was obtained (Zymoprep), and either electroporated into *E.coli* 10-beta electrocompetent cells (NEB) and further sequenced (Microsynth) or subjected to error-prone PCR (GeneMorph II Kit with 500 ng target DNA, 20 cycles) to introduce additional mutations.

Plasmids of desired dfRBP4 or RF2 variants were isolated and transformed into the *S. cerevisiae* strain EBY100 with the Frozen-EZ Yeast Transformation II Kit (Zymo Research) to individually characterize them. For *K*_D,_ and *EC*_50_ values based on yeast surface display, the geometric mean fluorescence intensity (gMFI) binding values were normalized and fitted to a 1:1 binding interaction model^41^. Negative values were set to 0. *T*_1/2_ values were calculated according to Zajc et al^41^.

### Mammalian cell culture

Buffy coats from healthy donors were purchased from the Austrian Red Cross, Vienna, Austria. Primary human T cells were purified by negative selection using the RosetteSep Human T cell Enrichment kit (STEMCELL Technologies) and cryopreserved in RPMI-1640 GlutaMAX medium (Thermo Scientific) containing 20 % FCS and 10 % DMSO (Sigma-Aldrich). T cells were cultivated between 0.3 - 2×10^6^ cells/mL in AIMV medium (Thermo Scientific) supplemented with 1 % glutamine (Gibco), 2.5 % Hepes (Pan Biotech), 2 % Octaplas (Blutspendezentrale Wien), and 200 U/mL recombinant human IL-2 (Peprotech). Raji cells (gift from Dr. Sabine Strehl, CCRI, Vienna) and Nalm-6 cells (ATCC), were transduced with a lentiviral vector encoding enhanced GFP_P2A_ffluc and maintained in RPMI-1640 GlutaMAX with 10 % FCS, 1 % penicillin–streptomycin (Thermo Scientific). Jurkat and Nur77 Jurkat reporter cells^33^ were cultivated in the same medium. Lenti-X 293T cells (Takara) were maintained in DMEM (Thermo Scientific) supplemented with 10 % FCS.

### *In vitro* transcription and mRNA electroporation

Transgenes were amplified via PCR and 200 ng of purified product were transcribed *in vitro* using the mMessage mMachine T7 Ultra Kit (Ambion) and the resulting mRNA was purified with the RNeasy Kit (Qiagen). Jurkat and Nur77 Jurkat cells were electroporated with 5 µg mRNA per construct using 4 mm cuvettes (VWR) and the Gene Pulser (Bio-Rad) with the following parameters: square wave protocol, 500 V, 3 ms.

### Intracellular protein complementation assay

Intracellular binding of proteins was determined with the Nano-Glo® Luciferase Assay System (Promega, live cell kit). The RBP4 and RF2 variants were N-terminally fused to an extracellular globular domain (truncated HER2 or truncated EGFR, respectively) to assess expression, and C-terminally fused to a subunit of the split Nanoluc (114 NanoBiT® “Small BiT” or 11S NanoBiT® “Large BiT” respectively). 16 h after co-electroporating the chains into Jurkat T cells, 100 000 cells were incubated in the presence of the appropriate compound (A1120, rapalog AP21967 or DMSO) for 30 min at 37 °C. The Nano-Glo reagents were reconstituted according to the manufacturer’s instructions and added to the cells immediately before measuring luminescence on an ENSPIRE Multimode plate reader (Perkin Elmer).

### Nur77 reporter cell assay

Expression of the CD19 antigen (SinoBiological, RefSeq BC006338) in Jurkat cells and of the CARs in Nur77 reporter cells was achieved via mRNA electroporation. Mock Nur77 cells were electroporated without mRNA as control. 18 h after electroporation, 25,000 effector Nur77 cells were co-cultured with Jurkat target cells expressing or not CD19 (E:T 1:2, 24 h at 37 °C) in RPMI-1640 GlutaMAX with 10 % FCS, 1 % penicillin– streptomycin containing the appropriate compound: 5 µM A1120, 0.5 µM AP21967 for FKBP/FRB dimerization, or medium only as control. mKO2 expression, caused by Nur77 activation, was measured with a BD FACSymphony A3 Cell Analyzer (BD Biosciences).

### Lentivirus production

The benchmark CD19 CAR and all split CAR chain II (CD3ζ – signaling) constructs were cloned into a third generation puromycin selectable pCDH expression vector (System Biosciences). Split CAR chain I (αCD19) constructs were cloned in a non-selectable pCDH vector with the puromycin gene removed and a stop codon introduced after the chain I.

Lenti-X 293 T cells (Takara) were co-transfected by second generation viral packaging plasmids pMD2.G and psPAX2 (Addgene plasmids #12259 and #12260, respectively; gifts from Didier Trono) and the desired expression vector with the PureFection Transfection Reagent (System Biosciences). Viral supernatants were collected on day 2 and 3 after transfection and concentrated 50- or 100-fold (for chain II or chain I, respectively) using the Lenti-X Concentrator (Takara). Viruses were resuspended in AIMV medium supplemented with 2 % Octaplas, 2.5 % Hepes, 1 % glutamine and 200 U/mL IL-2 and stored at −80 °C.

The benchmark CD20 CAR and the split transcription factors were cloned in a third-generation lentiviral transfer vector (Miltenyi Biotec). The benchmark CD20 CAR comprises an EF1α promoter driving a second-generation CAR with a Leu16-derived scFV, CD8 hinge and transmembrane domains, 4-1BB costimulatory domain and a CD3ζ activation domain. Split transcription factors were ordered as gBlocks (Integrated DNA Technologies) and cloned into the abovementioned transfer vector via conventional restriction-ligation method under the control of a PGK promoter.

Lentivirus were produced in adherent HEK 293T cells by co-transfection with transfer vector + helper plasmids (containing VSVG, gag-pol and rev) using polyethylenimine (PEI) as transfection reagent. Either 35 μg or 630 μg of DNA were transfected depending on the production scale (T175 flasks or 5-layer cell factories, respectively) in DMEM (Biowest) and incubated at 37°C for 6 hours before addition of 10% FCS. 10 mM sodium butyrate were added to the medium 24 hours post-transfection. Lentiviral particles were harvested 48- and 72-hours post-transfection by collection of the supernatant and filtration through a 0.45 µm hPES filter. Lentiviral particles were concentrated by centrifugation at 7000 g at 4°C overnight and resuspension of the viral pellet at 200X with TexMACS medium (Miltenyi Biotec) and stored at −80°C.

### CAR-T cell manufacturing via lentiviral transduction

For the generation of CD19 CAR and split CAR T cells, primary human T cells were thawed and activated on day 0 with Dynabeads Human T-Activator αCD3/αCD28 beads (Thermo Scientific) at a 1:1 bead-to-cell ratio. On day 1, 100 µL T cells (1 × 10^6^ cells/mL) were seeded in RetroNectin (Takara) coated 96 well plates and co-transduced with vectors encoding chain I and II by adding thawed viral supernatant at a 2:1:1 final dilution (cells:chain I:chain II). T cells expressing chain II were selected via puromycin selection (1 µg/mL, Sigma-Aldrich) on day 3.

The αCD19-containing chains (chain I of the split CARs or CD19 control CAR) include a FLAG-tag, the CD3z signaling chains (chain II) either have an N-terminal StrepII-tag (monomeric chains II) or are fused to mCherry (dimeric chains II), which were used to detect CAR expression and determine CAR^+^ fractions to set up proper E:T ratios.

iCD20 CAR T cells were generated by isolation of human primary T cells by magnetic-activated cell sorting (MACS®) using REAlease CD8 and CD4 MicroBead kit (Miltenyi Biotec), seeded at a density of 1 × 10^6^ cells/mL in TexMACS + premium grade human IL-7 + premium grade human IL-15 (Miltenyi Biotec) and activated with a 1:100 dilution of T Cell TransAct™ (Miltenyi Biotec). T cells were transduced with the first component of the transcriptional switch (D1.1 or D1.2) one day post-activation. Next day, cells were resuspended, spun down, supernatant was removed and the second component of the switch (D2.1 or D2.2) was transduced in TexMACS + premium grade human IL-7 + premium grade human IL-15 + 1:100 T Cell TransAct™ (Miltenyi Biotec). Frequency of transduced cells was determined via intracellular staining of DYKDDDDK (FLAG)-tag or HA-tag as indicated below. Frequency of induced CD20 CAR was detected by surface staining of the CAR using αCD20(Leu16) CAR antibody (Miltenyi Biotec).

### Intracellular immunostaining

Levels of the two different components of the split transcription factors were determined by intracellular staining of either DYKDDDDK (FLAG)-tag or HA-tag followed by flow cytometry. Cells were spun down by centrifugation at 300 g for 5 minutes, supernatant was removed, and cells were washed once with PBS. After spinning down and removing the PBS, cells were resuspended with ice-cold methanol and incubated 10 minutes at – 20°C. After two washes with PBS, cells were stained with either αDYKDDDDK (FLAG) antibody (APC, REAfinity 130-119-584, Miltenyi Biotec) or αHA-tag antibody (130-126-817, APC, Reaffinity, Miltenyi Biotec) at a dilution 1:50 in PBS + 2 mM EDTA 0.5% BSA following manufacturer’s protocol. After 15 minutes incubation at 4°C, cells were washed twice with PBS + 2 mM EDTA 0.5% BSA and resuspended in the same buffer for flow cytometric analysis using a MACSQuant 10 (Miltenyi Biotec). MACSQuantify 2.13.0 was used for data analysis.

### Flow cytometric analysis of CAR expression and cell counting

Nonspecific antibody binding was prevented by preincubating the cells for 10 min at 4 °C in FACS buffer (PBS with 0.2 % human albumin (CSL Behring) and 0.02 % sodium azide (Merck KGaA)) containing 10 % (v/v) human serum. Cells were then incubated with the antibodies (25 min at 4 °C) in FACS buffer, washed twice, and acquired on a BD FACSymphony A3 Cell Analyzer. Non-transduced or non-electroporated cells served as negative controls. The following antibodies were used: αFLAG (APC or PE; clone L5; BioLegend), αStrepII (FITC; clone 5A9F9; Genescript), αCD19 (APC or PE; clone HIB19, BioLegend), αCD3 (BV605; clone OKT3, Biolegend), αCD8 (BUV496; clone RPA-T8, BD Horizon), αCD4 (PerCP, clone OKT4; Biolegend), BD Horizon™ Fixable Viability Stain 780 (BD Biosciences).

When FACS panels included multiple brilliant violet-conjugated antibodies, all mixes and incubations were set up in brilliant stain buffer (BD Horizon). Serum-free washing buffer (plain PBS) was used when FACS panels included a fixable viability dye.

Accurate counts of cells in mice organs were obtained by adding 10 µL of AccuCheck Counting Beads (Thermo Scientific) to the cells after staining them and normalizing cell counts to the number of beads acquired.

### Cytotoxicity assays

All cytotoxicity assays were luciferase-based and carried out in RPMI without phenol red (Thermo Scientific) supplemented with 2 % Octaplas and 1 % Pen/Strep. Each experiment was set up with 3 technical replicates per condition. Raji and Nalm-6 target cells expressed enhanced GFP and luciferase.

CAR expression in primary human T cells was achieved via lentiviral transductions. CAR^+^ T cells were co-cultured with 10 000 target cells at an E:T ratio of 2:1 (37 °C, 6 h for Nalm-6 and 24 h for Raji target cells) in the presence of the appropriate compound. Mock T cells were spiked in to have the same total number of T cells among the different conditions and account for different CAR positive fractions. The next day luminescence was measured (ENSPIRE plate reader) and specific lysis was calculated according to the formula:

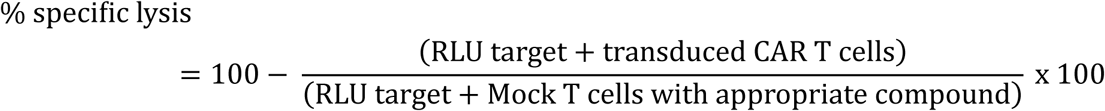

### Live-cell imaging-based cytotoxicity assays

Cytotoxic potential of iCD20 CAR systems was evaluated by live-cell imaging in co-cultures with CD20^+^ GFP^+^ 526-Mel tumor cells as previously described^42^. Briefly, either 1 × 10^4^ (for E:T 1:1) or 2 × 10^4^ (for E:T 1:2) CD20^+^ GFP^+^ 526-Mel cells per well were seeded in a 96-well plate with RPMI 1640 (Biowest) supplemented with 2 mM glutamine (Lonza) and 10% fetal bovine serum (FBS, Biochrome, Berlin, Germany) 16 hours before co-culture setup. Prior to each experiment, frequency of transduced cells among the different groups was adjusted by spiking-in non-transduced T cells in order to ensure equal number of transduced and total T cells in the co-culture. On the day of the co-culture, 1 × 10^4^ transduced T cells were added to the corresponding well of the 96 well plate containing attached CD20^+^ GFP^+^ 526-Mel cells. Co-cultures were incubated at 37°C + 5% CO_2_ in RPMI 1640 (Biowest) supplemented with 2 mM glutamine (Lonza) and 10% fetal bovine serum (FBS, Biochrome, Berlin, Germany) either in the absence or presence of 100 nM of A1120. Plates were monitored for 72 hours by live-cell imaging using an Incucyte S3 (IncuCyte) acquiring four pictures per well every two hours with an exposure time of 300 ms for the green channel. Green fluorescence integrated intensity of each at each time was calculated and normalized to time = 0 hours using Incucyte 2023A Rev2 software and exported for further analysis and graphical representation in GraphPad Prism.

### Cytokine release assays

Co-culture supernatants of triplicate wells were pooled and frozen for cytokine release assays. IFN-γ and IL-2 secretion was quantified by enzyme-linked immunosorbent assay (ELISA) with the ELISA MAX Deluxe Set Human IFN-γ and IL-2 respectively (Biolegend). The data was acquired on an ENSPIRE plate reader, and the analysis carried out with the Arigo GainData® online tool, excluding all values reaching the standard curve’s plateau.

For OFF kinetics, VAR-CAR T cells were cultivated with 5 µM A1120 72 h before the assay. At the indicated time points (48 h / 30 h / 24 h / 8 h / 4 h / 2 h prior to start of the co-culture assay), cells were washed to remove the drug and kept in culture in plain medium. A 24 h co-culture with Raji cells (E:T of 2:1) was then set up, followed by analysis of IFN-γ in the supernatant. VAR-CAR cells that were always in the presence of A1120 were washed once and A1120 was added (ON) or not (0 h) in the co-culture. Mock T cells and VAR-CAR T cells that were never cultured with A1120 (OFF) were used as negative controls. For ON kinetics, 5 µM A1120 was added in the VAR-CAR T cell cultures at the indicated time point (48 h / 24 h / 4 h / 2 h / 1 h) before the assay. CAR^+^ T cells were then co-cultured at an E:T ratio of 2:1 with Raji cells for 4 h (E:T of 2:1) and IFN-γ concentration in the supernatant was measured. For the 0 h time point A1120 was only added in the co-culture medium.

### *In vivo* experiments

NOD.Cg-Prkdc^scid^ Il2rg^tm1WJI^/SzJ (NSG, The Jackson Laboratory) mice were kept at the Core Facility Laboratory Animal Breeding and Husbandry of the Medical University of Vienna (CFL) under specific pathogen-free conditions according to FELASA recommendations (Himberg, Austria) or bred and kept at the IOV-IRCCS Specific Pathogen-Free animal facility (Padova, Italy). Animal experiments have been carried out either at the Institute for Molecular Pathology (IMP, Vienna) or Veneto Oncology Institute (IOV, Padova). All procedures were approved by the Magistratsabteilung 58, Vienna, Austria (GZ: MA 58 – 1390324-2024-19) and the Austrian Federal Ministry of Education, Science and Research or by the Italian Ministry of Health (Authorization n° 118/2020-PR), adhering to institutional guidelines (D.L. 26/2014). The study was conducted in compliance with the European Community Directive 86/609 for the care and use of laboratory animals and the FELASA and ARRIVE guidelines.

Primary human T cells were lentivirally transduced for CAR expression and expanded for 12 - 13 days.

For the VAR-CAR T cell experiment, cells expressing the CD19 CAR and the chain II of the VAR-CAR (CD3ζ containing, in pCDH vector) were selected via puromycin (1µg/mL) on day 3 after T cell thawing and activation. 0.5 × 10^6^ Raji cells expressing GFP and ffLuc were injected i.v. into the tail veins of NSG female mice (8 weeks old). Four days later, mice were randomized to ensure comparable mean tumor signals between groups, followed by i.v. injection of 4 × 10⁶ CAR-positive T cells.

For the iCD20 CAR stress test experiment, the cells were not selected. 0.5 × 10^6^ Raji cells expressing GFP and ffLuc were injected i.v. into the tail veins of NSG female mice (14 weeks old) and 0.5 × 10^6^ CAR positive T cells were injected i.v. five days later. In this case, Mock T cells were also injected into mice receiving the cCD20 CAR to account for very different CAR % and reach the same total number of T cells (20 Mio per mouse) compared to mice receiving iCD20 CAR Ts.

In both cases, A1120 was formulated in the chow^28^ by Research Diets (237 mg A1120 / kg chow, product C13695) and the same chow but not containing A1120 (product C13513, Research Diets) was used as negative control. Both chows were irradiated before use.

### *In vivo* bioluminescence imaging (BLI)

Bioluminescence imaging was performed at the IMP or IOV using an IVIS Spectrum imaging system (PerkinElmer). Mice were anesthetized with isoflurane (1000 mg/g Vetflurane, Virbac) and received i.p. injections of D-luciferin (150 mg/kg body weight, Goldbio). After 5 min, mice were transferred to the IVIS Imaging System and bioluminescence was measured with an automatic acquisition time in medium binning mode. Living Image^TM^ software (v.4.8, Revvity) was used to analyze the data and extract total photon fluxes. Pictures were cropped to reconstitute Figure 7D as groups were randomized based on tumor burden on d0 and mice in the same cage got different T cell treatments.

### Mouse organ processing

Blood was collected via tail veins or retro-orbital collection into EDTA tubes. Bone marrow single-cell suspensions were obtained by flushing two femurs with 15 mL PBS and subsequent filtering through a 70 µm cell strainer. Spleen samples were disrupted and passed through 70 µm filters twice. All single-cell suspensions were subjected to multiple washing steps in PBS and lysis of red blood cells with ACK Lysing Buffer. Finally, cells were blocked with 5 % Octaplas and a fraction of the volume was stained in 50 µL antibody mix (anti-hCD45-BUV395, 1:40; anti-hCD8-BV421, 1:20; anti-hCD3-BV510, 1:40; anti-hCD4-BV650, 1:40; anti-hCD19-PE, 1:10; anti-hCD62L-PE-Vio770, 1:50; anti-hCD45RA-AF647, 1:50; anti-TIM3-BV421, 1:50; anti-hLAG3-BV650, 1:50; anti-hCD39-PE-Vio770, 1:75; anti-CD3-BV605, 1:50; anti-Flag-PE 1:100; anti-hCD19-APC, 1:100; anti-StrepII-FITC, 1:25; fixable viability dye 780, 1:1000), washed twice and measured on a BD LSR Fortessa together with 10 µL AccuCheck Counting Beads (Thermo Scientific).

### Mass spectrometric analysis of A1120 concentrations in mice plasma

Whole blood was collected from mice in EDTA tubes and plasma was harvested by centrifuging at 2 000 g for 15 min and collecting the supernatant.

A1120 powder (Sigma Aldrich; PN: A3111) was used for the preparation of calibration standards. ^13^C_3_-labeled caffeine (Sigma Aldrich; PN: C082) served as the internal standard and was added in equal volumes to all calibration standards and samples. Calibration standards were spiked with negative plasma from control mice to achieve matrix matching. LC-MS grade acetonitrile (Thermo Scientific; PN: A956-1) was added to both standards and samples to precipitate plasma proteins. The mixture was cooled at 4 °C for 15 minutes, followed by centrifugation at 21 300 g for 15 minutes. The resulting supernatant was diluted with an equal volume of LC-MS grade methanol (Thermo Scientific; PN: A456-1) prior to injection.

Chromatographic separation was performed on a ZORBAX SB-Aq C18 column (Agilent, 2.1*50 mm, 1.8 µm) using 0.1 % formic acid in LC-MS grade water as the aqueous mobile phase (A) and 0.1% formic acid in 95 % LC-MS grade acetonitrile as the organic mobile phase (B). Initial conditions were set to 5 % B (0.5 min), ramped linearly to 80 % B over 4.5 min, held at 80 % B for 0.9 min, and then returned to baseline conditions for 2 min. The total method time was 8 minutes with a flow rate of 300 µL/min and a column oven temperature of 40 °C. The injection volume was 5 µL.

Detection was performed using an Orbitrap IQ-X Tribrid LC-MS^n^ mass spectrometer (Thermo Scientific) equipped with a H-ESI source in positive ion mode. The ion source parameters were set as follows: vaporizer temperature, 325 °C; ion transfer tube temperature, 275 °C; auxiliary gas pressure, 10 arbitrary units; sheath gas pressure, 50 arbitrary units; ion sweep gas pressure, 1 arbitrary unit; spray voltage, 3200 V. The Orbitrap resolution was set to 60 000, with an RF Lens setting of 60 % and a maximum injection time of 118 ms. RunStart EASY-IC was utilized for internal mass calibration. A tSIM method was employed for both A1120 ([M+H]^+^: 393.1421 *m/z*) and ^13^C_3_-labeled caffeine ([M+H]^+^: 198.0977 *m/z*). Data analysis was performed using TraceFinder 5.2.

### Statistical analysis

Statistical analyses were performed with GraphPad Prism software version 9 for Windows (GraphPad Software Inc.) and are indicated under each figure. *EC*_50_ values based on lysis or cytokine release were calculated with the ‘[Agonist] vs response function’ of GraphPad. *EC*_50_ values of inducible split transcription factors based on achieved frequencies and MFI of the induced gene of interest were calculated via non-linear fit of [Agonist] vs response model of GraphPad. Statistical analyses of cytokine secretion was performed on log-transformed values.

